# A Bayesian Approach for Inferring the Impact of a Discrete Character on Rates of Continuous-Character Evolution in the Presence of Background-Rate Variation

**DOI:** 10.1101/576207

**Authors:** Michael R. May, Brian R. Moore

## Abstract

Understanding how and why rates of character evolution vary across the Tree of Life is central to many evolutionary questions; *e.g.*, does the trophic apparatus (a set of continuous characters) evolve at a higher rate in fish lineages that dwell in reef versus non-reef habitats (a discrete character)? Existing approaches for inferring the relationship between a discrete character and rates of continuous-character evolution rely on comparing a null model (in which rates of continuous-character evolution are constant across lineages) to an alternative model (in which rates of continuous-character evolution depend on the state of the discrete character under consideration). However, these approaches are susceptible to a “straw-man” effect: the influence of the discrete character is inflated because the null model is extremely unrealistic. Here, we describe `MuSSCRat`, a Bayesian approach for inferring the impact of a discrete trait on rates of continuous-character evolution in the presence of alternative sources of rate variation (“background-rate variation”). We demonstrate by simulation that our method is able to reliably infer the degree of state-dependent rate variation, and show that ignoring background-rate variation leads to biased inferences regarding the degree of state-dependent rate variation in grunts (the fish group Haemulidae). [continuous-character evolution; discrete-character evolution; Bayesian phylogenetic comparative methods; data augmentation]

---

Variable rates of continuous-character evolution are central to many evolutionary questions. These questions may involve changes in the rate of character evolution over time (time-dependent scenarios) or among lineages (lineage-specific scenarios). Such questions may be pursued by means of agnostic surveys to detect rate variation (data-exploration approaches) or by testing predictions regarding factors hypothesized to influence rates of character evolution (hypothesis-testing approaches). A particular type of hypothesis posits that the rate of continuous-character evolution depends on the state of a discrete trait, *e.g.*, the evolutionary rate of the feeding apparatus (a set of continuous traits) in a lineage depends on the habitat type (the discrete trait) of its members.

The latter question—regarding state-dependent rates of continuous-character evolution—is currently pursued using a computational procedure (*e.g.*, Collar et al. 2009, 2010; Price et al. 2011, 2013) comprised of four steps: (1) fit a Brownian motion model to the observations (the tree and continuous-trait values at its tips), where the instantaneous rate of continuous-character evolution is assumed to be constant across all branches of the tree (the “null” or constant-rate model); (2) generate a sample of discrete-character histories (“stochastic maps”, Nielsen 2002; Huelsenbeck et al. 2003); (3) for each stochastic map, fit a Brownian motion model to the observations, where the instantaneous rate of continuous-character evolution at a given point on a given branch depends on the corresponding state of the discrete-character mapping (the “state-dependent model” O’Meara et al. 2006); (4) compare the fit of the state-dependent model (averaged over the sample of stochastic maps) to the constant-rate model using AIC. If the state-dependent model is preferred, we infer that rates of continuous-character evolution depend on the state of the discrete character.

The current approach has two potential problems. First, stochastic maps of the discrete character are generated without reference to the continuous characters. By construction, however, the state-dependent model specifies that the discrete and continuous characters are evolving *jointly*. The continuous characters therefore possess information about the history of the discrete character; disregarding this mutual information will lead to biased parameter estimates (Revell 2012a). Second, the null model—where the continuous characters are assumed to evolve at a constant rate across lineages—is extremely unrealistic. *Any* variation in the rate of continuous-character evolution—whether or not it is associated with the discrete character under consideration—is apt to be interpreted as evidence against the overly simplistic null model. This “straw-man effect” has the potential to mislead our inferences regarding the factors that impact rates of continuous-character evolution.

We describe a Bayesian approach for inferring the impact of a discrete trait on rates of continuous-character evolution that addresses the problems described above. We begin by developing a stochastic process that explicitly models the joint evolution of the discrete and continuous characters; this stochastic process can accommodate one or more continuous characters evolving under a state-dependent multivariate Brownian motion process. We refer to this new model as `MuSSCRat` (for Multiple State-Specific Rates of continuous-character evolution). We then develop an inference model that accommodates variation in the background rate of continuous-character evolution (*i.e.*, rate variation across lineages that is independent of the discrete character under consideration). This model of background-rate variation is spiritually similar to methods that agnostically survey for variation in rates of continuous-character evolution across lineages of a tree (*e.g.*, Eastman et al. 2011; Venditti et al. 2011). We implement this model in a Bayesian framework, which accommodates uncertainty in the phylogeny, discrete-character history, and parameters of the state-dependent model. We show by simulation that the method is able to reliably infer the state-dependent rates of continuous-character evolution, and that ignoring background-rate variation leads to an inflated false-positive rate. Finally, we demonstrate the new method with an empirical analysis of grunts (a group of haemulid fish) to illustrate the impacts of background-rate variation and prior specification on inferences about state-dependent rate variation.

## Methods

Our goal is to develop a state-dependent multivariate Brownian motion model, `MuSSCRat`, in a Bayesian statistical framework. We begin with a simple simulation to describe the parameters and basic properties of the `MuSSCRat` model. We then show how to calculate the probability of the process changing from one state to another over a fixed time interval under the `MuSSCRat` model (*i.e.*, how to compute transition probabilities). Next, we use these transition probabilities to compute the probability of observing the discrete and continuous characters across the tips of a phylogeny under the model (*i.e.*, how to compute the likelihood). Finally, we describe the relevant details—the priors and MCMC machinery—required to perform Bayesian inference under the `MuSSCRat` model.

### The State-Dependent Multivariate Brownian Motion Process

#### A simulation example

We introduce the salient properties of the `MuSSCRat` model by describing a simple simulation with a binary discrete character, *X*, and a single continuous character, *ϒ*, over a single branch. The discrete character has two states, 0 and 1; the continuous character can be any real number. The state of the simulation at time *t* is the pair of discrete- and continuous-character values, (*x*_*t*_, *y*_*t*_). (We use capital letters—*X* and *ϒ*—to represent random variables, and lowercase letters—*x* and *y*—to represent specific values of those random variables.)

The discrete trait evolves under a continuous-time Markov process, changing from state 0 to state 1 with rate *q*_01_, and from state 1 to state 0 with rate *q*_10_. The continuous character evolves under a state-dependent Brownian motion process, where the diffusion rate, 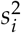, measures the rate of continuous-character evolution when the discrete character is in state *i* (*i.e.*, 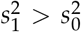 indicates that the con tinuous character evolves faster in discrete state 1 than in discrete state 0). In a small time interval of duration ∆*t*, where the discrete character begins in state *i*, the continuous character changes by a normally distributed random variable with mean 0 and variance 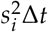, and the discrete character changes state with probability *q*_*ij*_Δ*t*.

We begin the simulation at time *t* = 0, with the discrete character in state *x*_0_ and the continuous character with value *y*_0_. We then increment the simulation forward in time by a small time interval, Δ*t*, applying the above rules describing how the state of the process changes during each time interval. We continue to increment the simulation forward in time until we reach the end of the branch (at time *T*). The outcome of the simulation is a *sample path* that records the state of the process from the beginning to the end of the branch (Fig. 1A).

**Figure 1:**
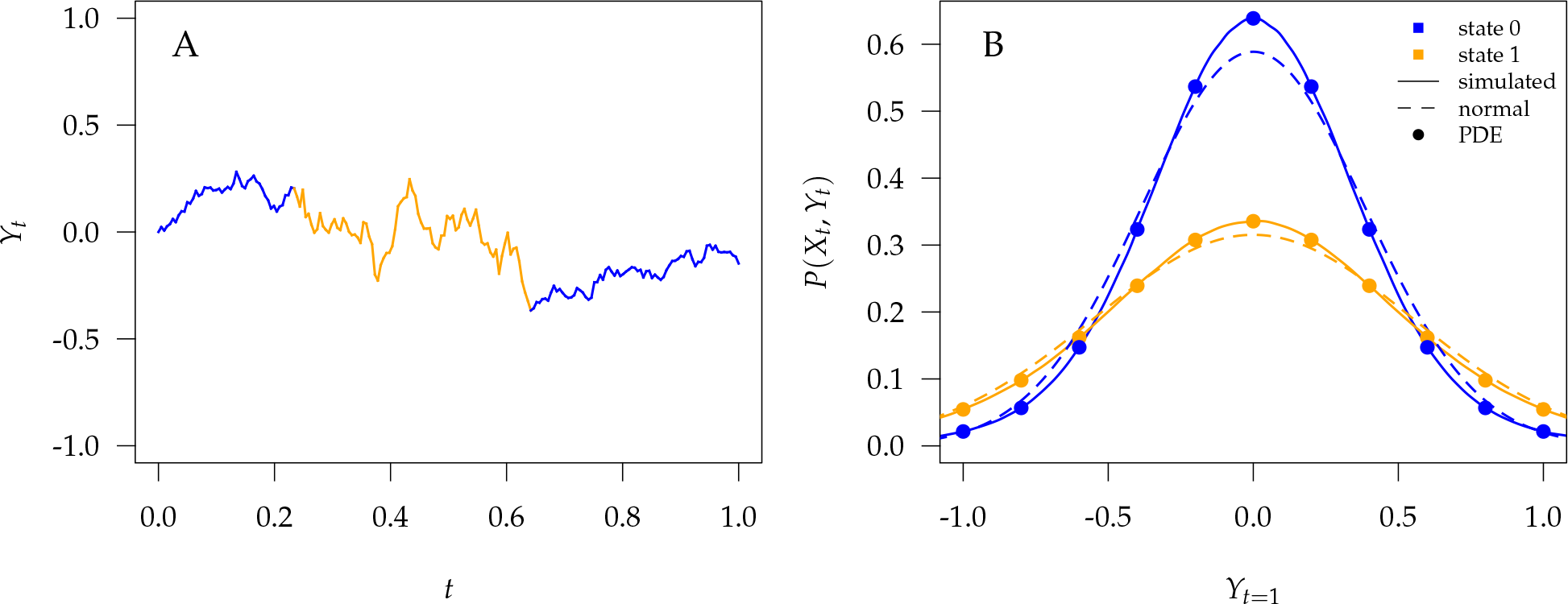
We perform simulations under the state-dependent multivariate Brownian motion process—the `MuSSCRat` model—to better understand its behavior. The process is either in discrete state 0 (blue) or discrete state 1 (orange), where the rate of change between discrete states is equal (*q*_01_ = *q*_10_ = 1), and the rate of continuous-character evolution is higher when the process is in the orange state 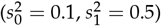. A) We simulate 0 1 a single sample path from *t* = 0 to *t* = 1. In this simulation, the process begins and ends in the blue state, but spends some time in the orange state. Note that there is more evolution in the orange state than in the blue state. B) We simulated 10 million sample paths and recorded their end states. Solid lines represent the simulated joint probability densities of the discrete and continuous states. Dashed lines represent the normal densities with parameters estimated from the simulated end states. Note that the simulated densities depart from the normal densities (both Kolmogorov–Smirnov *p*-value < 2E − 16). Colored points follow the densities computed by numerically solving the partial differential equations (1) and (2).

The *transition probability density* specifies the probability that the process ends in some state, (*x*_*t*_, *y*_*t*_), given an initial state (*x*_0_, *y*_0_), after a certain amount of time, *t*, has elapsed. We can approximate the transition probability for a given branch by simulating many sample paths and recording the end state for each simulation. The resulting frequency histogram of end states provides a Monte Carlo approximation of the transition probability density for a branch of duration *t*. Note that the transition probability densities of standard (*i.e.*, state-independent) Brownian motion processes are normal densities. By contrast, it is clear from our simulations that the transition probability densities under the state-dependent process are not normal densities (Fig. 1B). We describe two alternative approaches for calculating transition probability densities under the state-dependent process below.

#### The joint evolution of discrete and continuous characters

Our description of the state-dependent stochastic process implies that the relationship between discrete and continuous characters is unidirectional: *i.e.*, that the discrete trait influences the evolution of continuous characters, whereas the continuous characters do not influence the evolution of the discrete trait. Nevertheless, the continuous characters provide information about the history of the discrete character under the state-dependent process. Accordingly, it is potentially problematic to disregard the continuous characters when estimating the history of the discrete trait.

To illustrate the impact of the continuous characters on the discrete-character history, imagine that we evolve the process over a single branch of known duration, *T*. We assume that the process begins in discrete state 0 and ends in state 1, with a single change from 0 to 1 at some time, *t*. We also assume that the process begins in continuous state 0 and ends in continuous state *y*_*T*_ (which we observe). Finally, we assume that the continuous character evolves at a higher rate in discrete state 1 than in discrete state 0 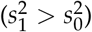.

It is clear that the amount of continuous-character evolution over the branch depends on the timing of the discrete-character change: an earlier change to state 1 provides more opportunity for the continuous character to evolve. Therefore, if we observe that the continuous character has changed substantially along the branch, it is more probable that the change to state 1 occurred earlier. In other words, the observed value of *y*_*T*_ influences the probability of the history of the discrete trait (Fig. 2).

**Figure 2:**
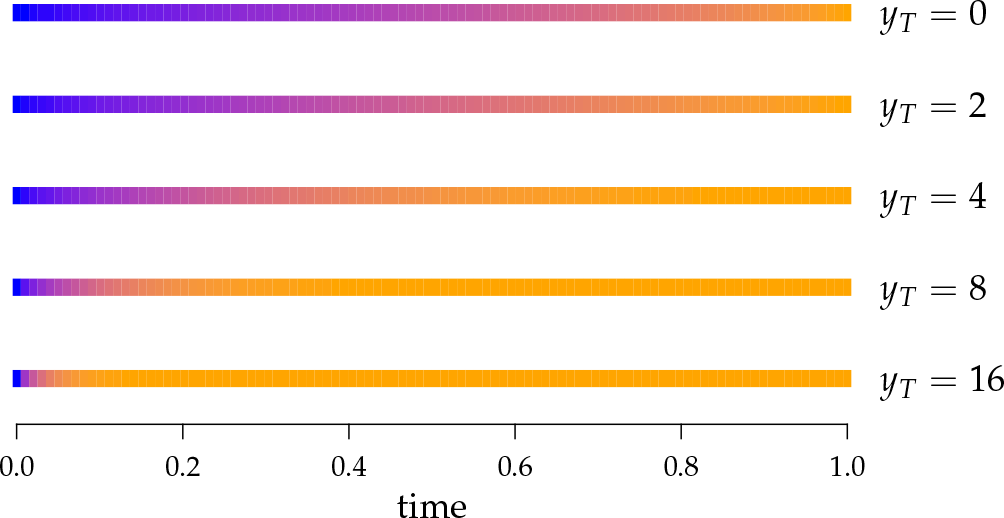
The probability of the discrete-character history depends on the observed continuous character. Here, we assume that the discrete trait begins in state 0 and ends in state 1, with exactly one 0 → 1 change. We also assume that the continuous character begins with value 0, and evolves at a rate that depends on the state of the discrete trait; 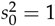 and 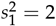. Each row corresponds to a different end state for the continuous character, *y*_*T*_, with larger values of *y*_*T*_ implying more evolution. Blue and orange shades indicate a high probability of being in discrete state 0 and 1, respectively. Larger values of *y*_*T*_ require more continuous-character evolution, implying that the discrete trait changed earlier in its history.

The conventional procedure for detecting state-specific rates of continuous-character evolution (*e.g.*, Collar et al. 2009) treats the history of the discrete character as independent of the observed continuous characters; *i.e.*, it generates stochastic maps of the discrete character without reference to the continuous characters, fits a continuous-character model to each simulated discrete-character history, and then averages parameter estimates over the sample of discrete-character histories *uniformly*. Accordingly, this approach ignores the impact of the continuous-character data on the distribution of discrete-character histories. The approach that we pursue here explicitly models the joint evolution of the discrete and continuous characters.

#### Parameters of the MuSSCRat model

We model the joint evolution of a discrete binary character, *X*, and a set of *c* continuous characters, ***ϒ***, as a stochastic process. The discrete trait has two possible values, which we arbitrarily label 0 and 1; *x* ∈ (0, 1). The continuous characters are a vector of real-valued random variables, where *y*[*i*] is the value of the *i*^th^ continuous character 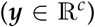. Variables of the `MuSSCRat` model are summarized in Table 1.

**Table 1:** The variables of the `MuSSCRat` model and their interpretation.

| Variable | Interpretation |
| --- | --- |
| $Y$ | The continuous characters for one lineage |
| $y_t$ | The state of the continuous characters at time $t$ |
| $c$ | The number of continuous characters |
| $\mathcal{Y}$ | An $n \times c$ matrix containing the $c$ continuous characters for the $n$ species in the tree |
| $X$ | The discrete character for one lineage |
| $x_t$ | The state of the discrete character at time $t$ |
| $\mathcal{X}$ | An $n \times 1$ matrix containing the discrete character for the $n$ species in the tree |
| $\kappa_l$ | The complete history of the discrete character (the state at the beginning and end of the process, and all of the changes in between) along the lineage $l$ |
| $\tau(\kappa_l, i)$ | The amount of time that the discrete-character history $\kappa_l$ spent in discrete state $i$ |
| $Q$ | The instantaneous-rate matrix of the discrete-character CTMC model |
| $q_{ij}$ | The instantaneous rate of change from state $i$ to state $j$ |
| $\beta^2$ | The background rate of continuous-character evolution among all lineages |
| $\beta_l^2$ | The background rate of continuous-character evolution for lineage $l$ |
| $\sigma^2$ | The relative rates of continuous-character evolution for each continuous character |
| $\zeta^2$ | The relative rates of continuous-character evolution for each discrete state |
| $R$ | The evolutionary correlation matrix |
| $\rho_{ij}$ | The evolutionary correlation between continuous characters $i$ and $j$ |
| $\Sigma$ | The discrete-state-independent evolutionary variance-covariance matrix, $\Sigma = \sigma R \sigma$ |
| $\Sigma^i$ | The evolutionary variance-covariance matrix for discrete-state $i$, $\Sigma^i = \zeta_i^2 \Sigma$ |
| $\Sigma_l$ | The evolutionary variance-covariance matrix for lineage $l$, $\Sigma_l = \beta_l^2 [\tau(\kappa_l, 0)\Sigma^0 + \tau(\kappa_l, 1)\Sigma^1]$ |

We assume that the discrete character evolves under a continuous-time Markov process, and that the continuous characters evolve under a multivariate Brownian motion process with rates that depend on the state of the binary character. These model components collectively describe how the set of characters, (*x*, ***y***), evolve together over a single branch; we detail the evolutionary dynamics of this process over an entire tree when we describe the likelihood function.

The instantaneous-rate matrix, *Q*, describes the rates at which the binary character evolves: *q*_*ij*_ describes the instantaneous rate of change from discrete state *i* to discrete state *j*:

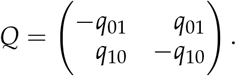

The complete history of the discrete trait on branch *l*—which we represent as *κ*_*l*_—specifies the state of the character at the beginning and end of the branch, and also the times of any character-state changes along the branch. We refer to the amount of time the history spends in state *i* as *τ*(*κ*_*l*_, *i*).

We assume the continuous character evolves under a multivariate Brownian motion model with some (scalar) base rate, 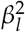 (the subscript refers to a specific branch, *l*). While in discrete state *i*, the base rate is multiplied by the state-specific relative rate, 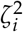. We also allow the relative rate of evolution to vary among the continuous characters. The vector ***σ***^**2**^ contains these relative rates; 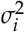 is the relative rate of continuous character *i*. The evolutionary correlations between characters are contained in the *c* × *c* symmetric correlation matrix, *R*, where [*R*]_*ij*_ = *ρ*_*ij*_ specifies the correlation between characters *i* and *j*.

We assume that the relative rates of change between characters, ***σ***^**2**^, and the evolutionary correlations between characters, *R*, are independent of the discrete state; in other words, we assume that the state of the discrete trait affects only the *overall rate* of continuous-character evolution, but not the *nature* of the evolutionary process (as represented by ***σ***^**2**^ and *R*). We combine the relative rates among characters and the correlation matrix to form the overall evolutionary variance-covariance matrix, Σ:

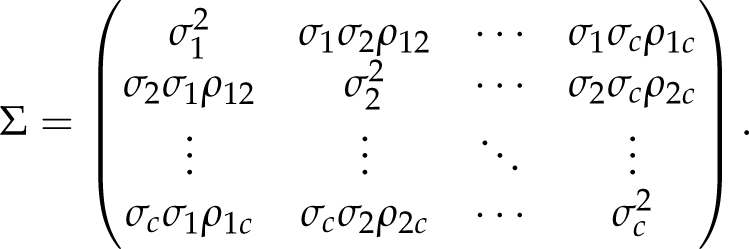

Similarly, we represent the state-dependent evolutionary variance-covariance while the process is in state *i* as Σ^*i*^:

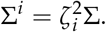

Finally, we represent the evolutionary variance-covariance matrix for branch *l* with discrete-character history *κ*_*l*_ as Σ_*l*_:

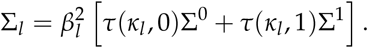

### Transition Probabilities

The transition probability density specifies the probability that the process changes from state (*x*_*i*_, ***y***_*i*_) to state (*x*_*j*_, ***y***_*j*_) over an interval of time *t* under the state-dependent stochastic process. Transition probabilities are central to computing the probability of realizing the observed character data under the `MuSSCRat` model (*i.e.*, the likelihood, described below).

In this section, we first describe how to compute the “full” transition probability density of the state-dependent process; this involves simultaneously integrating over all histories of changes in both the discrete and continuous characters. We then describe how to compute the “conditional” transition probability density; this involves integrating over all histories of the continuous characters for a given history of the discrete character. Finally, we demonstrate how numerically integrating the conditional transition probability density over discrete-character histories provides a theoretically valid—and computationally tractable—approximation to the full transition probability density.

#### Computing the full transition probability densities

The full transition probability density is the probability that the process changes from an initial state, (*x*_*i*_, ***y***_*i*_), to an end state, (*x*_*j*_, ***y***_*j*_), after a specified interval of time, *t*: *P*(*X*_*t*_ = *x*_*j*_, ***ϒ***_*t*_ = ***y***_*j*_ | *X*_0_ = *x*_*i*_, ***ϒ***_0_ = ***y***_*i*_, *θ*, *t*), where *θ* contains all the parameters of the `MuSSCRat` model. First, consider that the probability that the process is in discrete state *i* and continuous state ***y*** at time *t* is *P*_*i*_(***y***, *t*); for example, the densities described by the solid curves in Figure 1B correspond to *P_i_*(***y***, *t*).

In principle, the transition probability density can be calculated by solving a set of partial differential equations (derived in the Supplemental Material):

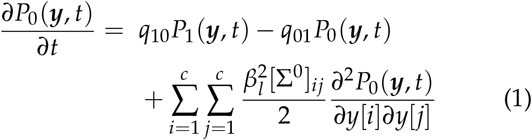

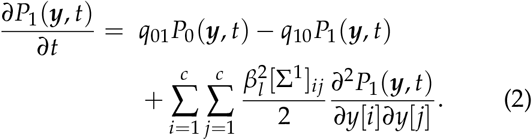

The first two terms on the right-hand-side of each equation represent the flow of the probability into and out of the discrete states, respectively, under a continuous-time Markov process; the third term represents the change in the probability density of the continuous character, which is a function of the discrete state.

Together, these PDEs describe the rate of change of the probability density, *P*_*i*_(***y***, *t*). Accordingly, if we knew the initial state of the process, we could use these equations to compute the probability that the process is in a given state, *P*_*i*_(***y***, *t*), at some later time, *t* (*i.e.*, we could compute the transition probability density). Unfortunately, it is not clear that these PDEs can be solved analytically, which motivates our development of an approach based on numerical integration of conditional probability densities (described below).

#### Computing the conditional transition probability densities

An alternative to solving the partial differential equations described above is based on assuming the complete history of the discrete character on branch *l*, represented by the variable *κ*_*l*_, is known. In this case, the PDE describing the rate of change of the probability density of the continuous characters while in a particular discrete state is:

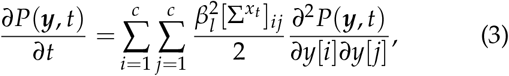

where 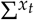 is the state-dependent variance-covariance matrix corresponding to the (assumed) discrete-state at time *t*.

Assuming that the process is in discrete state *i* for duration *t*_*i*_, this PDE has a known analytical solution: the transition probability density is a multivariate-normal distribution with mean ***y***_0_ and variance-covariance matrix 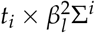. This implies that the changes in ***y*** while in discrete state *i*, Δ_*i*_, are multivariate-normally distributed:

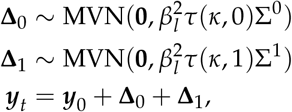

where *τ*(*κ*_*l*_, *i*) is the amount of time the history spends in discrete state *i* and **0** is a 1 × *c* vector of zeros (indicating that the expected amount of change for each continuous character is 0). Because it is the sum of multivariate-normally distributed random variables, ***y***_*t*_ is also multivariate-normally distributed:

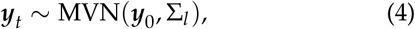

where 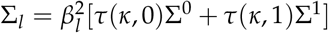 is the branch-specific variance-covariance matrix given the discrete character history and the base-rate of evolution, 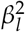. The conditional transition probability density for a given discrete-character history is therefore the familiar multivariate-normal probability density function:

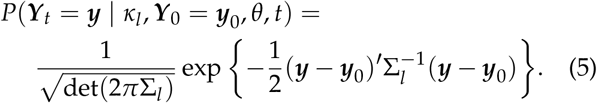

We can compute the *full* transition probability density by integrating the conditional transition probability density over all possible discrete-character histories, *K*, in proportion to their probability under the continuoustime Markov process:

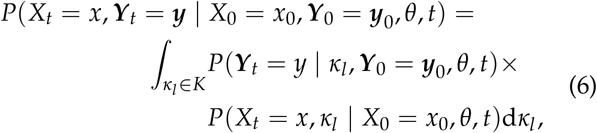

where *P*(*X*_*t*_ = *x*, *κ*_*l*_ | *X*_0_ = *x*_0_, *θ*, *t*) is the joint probability of the end state and the assumed character history of the discrete character:

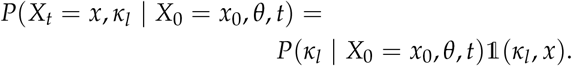

The probability of the assumed discrete-character history, *P*(*κ*_*l*_ *X*_0_ = *x*_0_, *θ*, *t*), can be computed by exploiting the fact that waiting times between discrete-character changes are exponentially distributed (Fig. 3), and 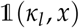, *x*) is an indicator function that evaluates whether the end state of the discrete-character history is equal to *x*:

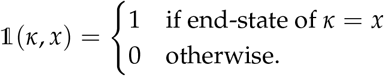

**Figure 3:**
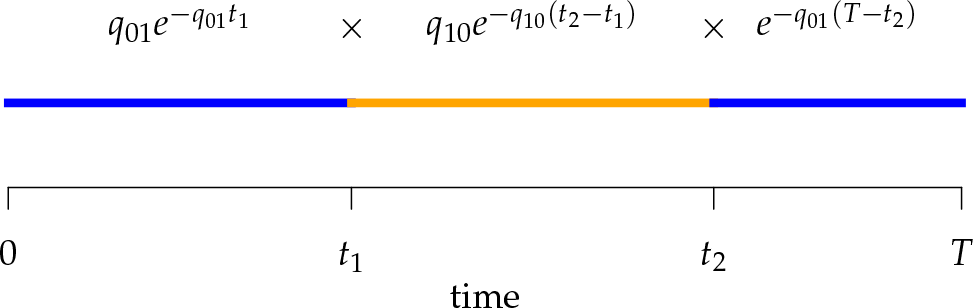
Computing the probability of a character history, *κ*_*l*_. We simulate the process forward in time with rates specified in the matrix *Q*. Blue and orange segments correspond to discrete states 0 and 1, respectively. The probability of the history is the product of the probabilities of waiting times between events (or the probability of no event in the final segment) given the current rate of change.

Equation (6) makes it clear that numerically integrating the conditional transition probability density over the discrete-character histories provides a theoretically valid approximation to the full transition probability density. There are many numerical-integration algorithms to choose from; a familiar option is Monte Carlo integration. First, we simulate *n* discrete-character histories, 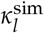, under the specified *Q* matrix, then we average the conditional transition probability density over the set of simulated discrete-character histories to approximate the full transition probability density:

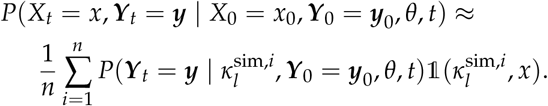

The approach that we develop uses a spiritually similar numerical-integration strategy—Markov chain Monte Carlo integration with data augmentation—to estimate the posterior distribution of parameters in the Bayesian model (described below).

### Likelihood

The likelihood function allows us to compute the probability of realizing the observed discrete and continuous characters at the tips of a phylogeny under the `MuSSCRat` model. In this section, we first describe the “full” likelihood function, which relies on combining the *full* transition probability densities (described previously) for all branches of the tree. We then describe the “augmented” likelihood function, which relies on combining the *conditional* transition probability densities (described previously) for all branches of the tree. The augmented likelihood allows us to efficiently calculate the joint probability of the discrete and continuous characters *conditional* on an assumed discrete-character history. Our Bayesian implementation of the `MuSSCRat` model uses a Markov chain Monte Carlo algorithm—*data augmentation*—to numerically integrate the augmented likelihood over all possible discrete-character histories (described below).

We imagine that we have sampled one discrete character and *c* continuous characters for each of *n* species; relationships among these species are defined by the phylogeny, Ψ. We store the discrete characters in an *n* × 1 column vector, 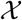, and the continuous characters in an *n* × *c* matrix, 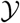. We assume that the discrete and continuous characters evolve independently along each of the 2*n* − 2 branches of the phylogeny. We index the internal nodes according to their sequence in a post-order traversal of the tree, starting from the root (which has index 1).

#### The full likelihood

Our goal is to compute the joint probability of the character data, 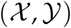, given the model parameters and the phylogeny, 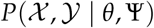, where *θ* contains all the parameters of the `MuSSCRat` model. To aid the calculation, we imagine that we also know the state of the discrete character and the values for all continuous characters at each internal node of the tree. We index the discrete state and the continuous states at the beginning of branch *l* as *x*_*b*(*l*)_ and *y*_*b*(*l*)_, respectively. Similarly, we index the discrete and the continuous states at the end of branch *l* as *x*_*e*(*l*)_ and *y*_*e*(*l*)_, respectively. Accordingly, the transition probability density for branch *l* is simply the transition probability density for the corresponding initial and end states of that branch; *P*(*x*_*e*(*l*)_, *y*_*e*(*l*)_ | *x*_*b*(*l*)_, *y*_*b*(*l*)_, *θ*, *t*_*l*_), where *t*_*l*_ is the duration of branch *l*. The probability of the states at the root of the tree is *P*(*x*_1_, ***y***_1_ | *θ*). We combine the transition probabilities for all branches as their product (because the process evolves independently along each branch),and average over the states at the internal nodes and root. This provides the joint probability of the character data (observed at the tips of the tree) under the state-specific model (equation [7]). We note that each integral over ***y***_*i*_is in fact a multidimensional integral over 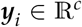.

#### The augmented likelihood

As noted previously, we do not know an analytical solution to the transition probability density along a branch, which makes it difficult to evaluate equation (7). We circumvent this issue by *augmenting* the observed data with the complete history of the discrete character along each branch of the tree. We include a vector of character histories, ***κ***, where *κ*_*l*_ is the discrete-character history along branch *l* (including the state at the beginning and end of the branch).

Conditioning on the discrete-character history provides two benefits: (1) it allows us to ignore the summation over discrete-character states at internal nodes in equation (7), and; (2) it allows us to use the *conditional transition probability density*, which follows a known multivariate normal density (equation [5]). The *augmented likelihood* is a product of the joint probability of 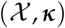 and the conditional probability density of 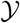 given ***κ*** (equations [8] and [9]). The full likelihood can be computed by integrating the augmented likelihood over all discrete-character histories:

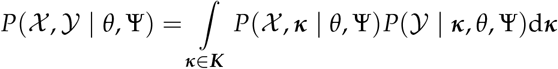

We compute the joint probability of 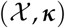 as a product of independent probabilities across each of the 2*n* − 2 branches:

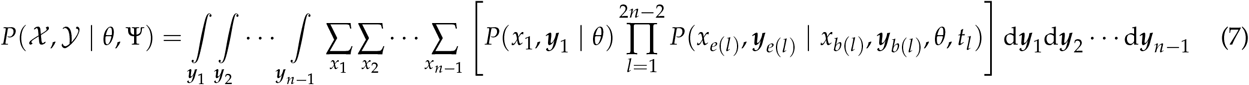

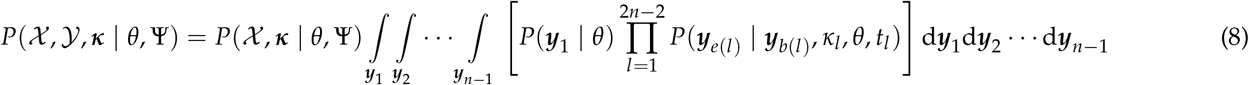

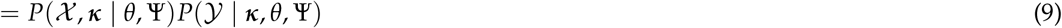

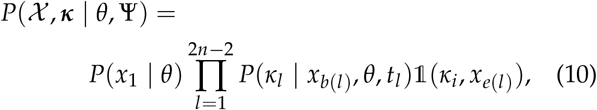

where *x*_*e*(*l*)_ refers to *observed* values of *X* for terminal branches, while the remaining *x*_*b*(·)_ and *x*_*e*(·)_ refer to *augmented* values of *X* at internal nodes. We compute *P*(*κ*_*l*_ | *x*_*b*(*l*)_, *θ*, *t*_*l*_) as illustrated in Figure 3. By convention, we use the stationary distribution of the Markov chain defined by *Q* as the probability of the root state, *P*(*x*_1_ | *θ*). We can compute the conditional probability density of the continuous characters, 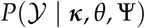, using standard algorithms to integrate over the distributions of the states at internal nodes. Specifically, we use Felsenstein’s REML algorithm (Felsenstein 1973, 2004), extended to multivariate Brownian motion (Huelsenbeck and Rannala 2003; Freckleton 2012), to compute the conditional probability density of 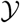. This algorithm assumes a uniform prior over all possible continuous states at the root.

#### Accommodating background-rate variation

In reality, the rate of continuous-character evolution may vary across lineages for reasons other than the discrete character of interest; we call this “background-rate variation”. Failing to accommodate background-rate variation may mislead our inferences about the “foreground” parameter, ***ζ***^**2**^. We can relax the assumption that all branches share a common background rate by allowing the background-rate parameter, 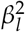, to vary among branches. The “constant-background-rate model” is therefore a special case of the background-rate-variation model; *i.e.*, when 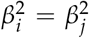 for all *i*, *j* branches.

Incorporating background-rate variation in the `MuSSCRat` model does not complicate the computation of the augmented likelihood, since it simply “rescales” the variance-covariance matrices on each branch. However, including background-rate variation *does* complicate inference; specifically, it causes the `MuSSCRat` model to become nonidentifiable (see Supplemental Material). A model is nonidentifiable when multiple combinations of parameters have identical likelihoods (Rannala 2002; Ponciano et al. 2012). Consequently, parameters of a nonidentifiable model cannot be estimated by standard maximum-likelihood methods because there may be no unique “maximum” likelihood. In this case, it is necessary to apply constraints on nonidentifiable parameters that “penalize” different combinations of parameters that have identical likelihoods. Bayesian models provide a natural solution to nonidentifiability, as the prior distributions on the parameters naturally penalize combinations of parameters that might have identical likelihoods (that is, the joint posterior probability of parameter combinations with identical likelihoods will differ if their joint prior probabilities are different). This motivates our development of a Bayesian model for inferring parameters of the `MuSSCRat` model with background-rate variation.

### Bayesian Inference

We implement the `MuSSCRat` model with backgroundrate variation as a Bayesian model to infer the joint posterior density of the model parameters given the observed data, 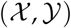, and the assumed tree, Ψ. The joint posterior density represents our beliefs about the parameter values after observing the data; it is the product of the likelihood—the probability of observing the data given the model parameters—times the joint prior density (reflecting our beliefs about the parameter values before observing the data). We must therefore specify both the likelihood function and also the joint prior density to compute the joint posterior distribution.

We use the augmented likelihood function (described in the previous section), and include the character histories along each branch, ***κ***, as variables in the model. In effect, we are “augmenting” the discrete-character data observed at the tips of the tree with unobserved discrete-character histories over the entire tree; this technique is referred to as *data augmentation* (Tanner and Wong 1987; Robinson et al. 2003; Mateiu and Rannala 2006; Landis et al. 2013).

We use a joint prior density comprised of the prior densities for each of the model parameters, described below. Having specified the augmented likelihood function and the joint prior density, we can write down the joint posterior density of the model parameters and the discrete-character histories:

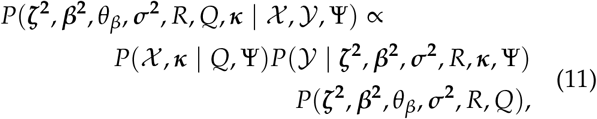

where the first two terms on the right-hand-side are the augmented likelihood, the third term is the joint prior density on the model parameters, and *θ*_*β*_ are the parameters of the background-rate-variation model.

The denominator in equation (11)—the *marginal likelihood* (not shown)—is a multidimensional integral of the augmented likelihood function over the joint prior density. The marginal likelihood is impossible to compute; the joint posterior probability density is therefore also impossible to solve analytically. Therefore, we approximate the joint posterior density using a numerical algorithm, Markov chain Monte Carlo. In addition to approximating the joint posterior density of the model parameters, our MCMC algorithm includes proposals that serve to numerically integrate over all possible discrete-character histories in proportion to their probability under the full state-dependent process.

The above Bayesian model conditions on a known tree, Ψ. In principle, it is easy to relax the assumption of a fixed tree by including it as a parameter in the model. In this case, we may include a sequence alignment, 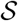, to improve our ability to estimate the topology and branch lengths/divergence times. The joint posterior distribution (equation [11]) becomes:

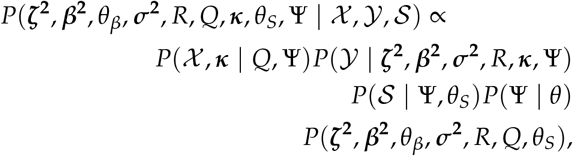

where *θ*_*S*_ contains all of the parameters of the phylogenetic model.

### Priors

We assume that the background-rate parameters, ***β***^**2**^, are drawn from a hierarchical model with parameters *θ*_*β*_, and the remainder of the parameters are 10drawn from independent prior distributions, so that the joint prior density becomes:

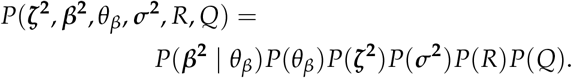

We describe our prior distributions in the following paragraphs; these parameterizations reflect our “baseline” model, but we explore alternative priors and prior sensitivity in our empirical analyses.

We draw the lineage-specific background rates of continuous-character evolution, ***β***^**2**^, iid from a shared lognormal distribution with mean *μ* and standard deviation *ν*. We use a uniform prior on log_10_(*μ*), such that *μ* is drawn from a log_10_-uniform distribution between 10^*−*3^ and 10. We draw the standard deviation, *ν*, from an exponential distribution with mean *H*. The constant *H* is the standard deviation for a lognormal distribution that indicates that our 95% prior belief ranges over one order of magnitude (see Supplemental Material). This model is the continuous-character analog of the uncorrelated lognormal (UCLN) relaxed-clock model used to describe variation in rates of molecular evolution across lineages (*e.g.*, Drummond et al. 2006; Lemey et al. 2010). Accordingly, we refer to this extension of the `MuSSCRat` model with background-rate variation as the `MuSSCRat` + UCLN model. A convenient property of the UCLN model is that—as *ν* shrinks to 0—it collapses to a “strict” morphological-clock model, where 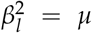 for all lineages. Our prior on *ν* specifies that we expect the values of ***β***^**2**^ to range over about one order of magnitude, but the exponential prior allows the standard deviation to shrink to 0 if the data prefer a strict morphological clock. In summary, we specify the background-rate-variation component of the prior model as:

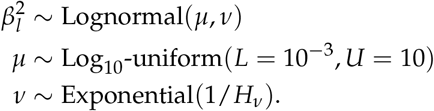

The parameter vector ***ζ***^**2**^ describes the relative rate of continuous-character evolution for each of the discrete states. We specify a Dirichlet distribution on *half* the values of ***ζ***^**2**^. Specifying the prior on half the values of ***ζ***^**2**^ ensures that the mean value of ***ζ***^**2**^ is 1, which allows us to interpret these parameters as the *relative* rate of continuous-character evolution in the alternative discrete states. We assume the concentration parameters of the Dirichlet distribution are the same, so this is a symmetric Dirichlet distribution with parameter 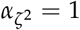:

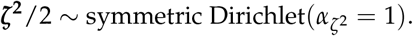

The average rate of change for each of the *c* continuous characters may vary; we allow the relative rate of continuous characters to vary by including a parameter vector, ***σ***^**2**^, where 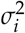 is the rate of evolution of the *i*^th^ continuous character. We specify a Dirichlet distribution on 1/*c*^th^ of the values of ***σ***^**2**^. We adopt the same logic as above for the prior on ***ζ***^**2**^, specifying a symmetric Dirichlet distribution with parameter 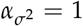 such that the mean value of ***σ***^**2**^ is 1:

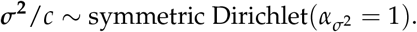

The symmetric matrix *R* determines the evolutionary correlation between each pair of continuous characters; *ρ*_*ij*_ = *ρ*_*ji*_ is the correlation between characters *i* and *j*. The matrix *R* has a special constraint that is must be positivesemidefinite, which makes it difficult to specify, for example, iid priors on each *ρ*_*ij*_. We use the LKJ distribution as a prior on *R*, which defines a prior over positive-semidefinite correlation matrices (Lewandowski et al. 2009). Briefly, this distribution draws *partial correlations*, *p*_*ij*_, from a sequence of Beta distributions such that the induced correlations, *ρ*_*ij*_, have the same marginal densities that are centered on 0. The resulting correlation matrices have prior density:

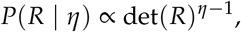

where *η* > 0 is inversely related to the variance of the correlation parameters: larger values of *η* result in marginal distributions on *ρ*_*ij*_ that are concentrated closer to 0, while smaller values of *η* result in distributions that are more diffuse. We choose *η* = 1, which indicates a uniform distribution over all possible positive-semidefinite correlation matrices:

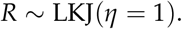

Finally, the matrix *Q* describes the rates of change between the discrete-character states. We note that the matrix *Q* has stationary frequencies, ***π***:

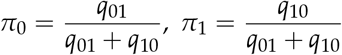

and that, at stationarity, the expected number of transitions, *k*, over an interval of duration *t* is:

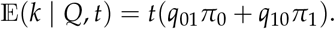

Assuming that the rates of change are symmetric (*q*_01_ = *q*_10_) or asymmetric (*q*_01_ ≠ *q*_10_) may have some impact on our analysis through the distribution on ***κ***. Moreover, inferring whether rates of change are (a)symmetric is often of direct interest to researchers studying discrete-character evolution. We therefore specify a mixture distribution on ***q***, so that ***q*** may be symmetric or asymmetric. Specifically, we draw ***q***, from a degenerate distribution concentrated on equal rates, 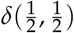, with probability *p* = 0.5, and from a Dirichlet distribution with parameter *α*_*q*_ = 1 with probability 1 − *p* = 0.5.

We place the rates, ***q***, in a matrix *Q*′, rescaled so that the mean rate of change, *r*, is 1:

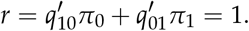

We include an additional parameter, *λ*, which scales the overall rate of change, so that:

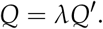

Constructing the matrix *Q* in this way means that the expected number of transitions over an interval of duration *T* is 𝔼(*k* | *Q*, *T*) = *λT*. We draw *λ* from a lognormal prior distribution with standard deviation *H*, and specify the mean such that the expected number of transitions over the entire phylogeny is *k*. The prior expected number of transitions reflects an empirical Bayesian prior, and should be specified differently for different datasets; for the simulations and analyses we describe later, we use 𝔼(*k* | *Q*, *T*) = 5 (where *T* is the sum of all branch durations in the tree). In summary, our overall prior on *Q* is:

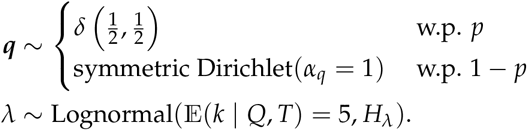

#### Constant background rates

The `MuSSCRat` model with constant-background rates is nested within the model with background-rate variation described above: as *ν* → 0, the lognormal prior on ***β***^**2**^ collapses to a point centered on *μ* (so that all values of ***β***^**2**^ become increasingly similar).

To specify the constant-background-rate model explicitly, we draw a single value for *β*^2^ from a log_10_-uniform distribution between 10^−3^ and 10. Otherwise, we use the same prior distributions for the constant-background-rate model as we described for the variable-background-rate model, above. The resulting posterior distribution is:

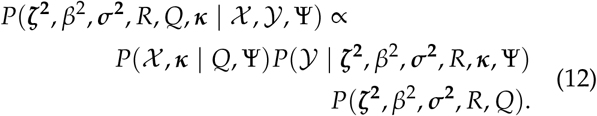

#### Markov chain Monte Carlo

The joint posterior probability density cannot be calculated analytically because we cannot evaluate the marginal likelihood. We therefore approximate the joint posterior probability density numerically using Markov chain Monte Carlo (MCMC); specifically, we draw samples from the joint posterior distribution using the Metropolis-Hastings and Green algorithms (Metropolis et al. 1953; Hastings 1970; Green 1995). We use standard proposal distributions for the majority of the parameters in our model; for brevity, we only provide details for two of our more uncommon proposal distributions—for moves between symmetric and asymmetric *Q* matrices, and for the discrete-character histories—in the Supplemental Material.

Our data-augmentation strategy involves including the complete history of the discrete character, ***κ***, as a variable in the Markov chain. As such, the MCMC procedure includes proposals that change the discrete-character history. When a new character history, ***κ′***, is proposed, it is accepted with probability *A*, computed as:

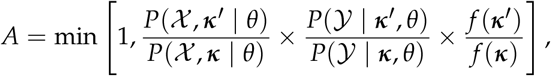

where *f* (***κ***) is the distribution from which the new character history is drawn. Note that the probabilities of the discrete characters, 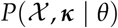, and continuous characters, 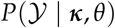, both contribute to the probability that the proposed discrete-character history is accepted. Importantly, this means that the continuous characters are able to correctly influence the discrete-character histories; *i.e.*, we are correctly modeling the joint distribution of the discrete and continuous characters.

#### Implementation

We implemented our `MuSSCRat` model in the open-source Bayesian phylogenetic software, `RevBayes` (Höhna et al. 2016). Our implementation relies upon the discrete-data augmentation functionality developed in `RevBayes` by Michael J. Landis and Sebastian Höhna (unpublished), extended to accommodate phylogenetic uncertainty. Owing to the flexibility of `RevBayes`, our implementation allows users to explore the impact of binary or multistate discrete traits on rates of continuous-character evolution, provides tremendous flexibility for specifying priors, enables simultaneous inference of ancestral states for both discrete and continuous characters, and allows joint inference of the phylogeny, divergence times, and parameters of the `MuSSCRat` model. We provide `Rev` scripts for performing analyses under the `MuSSCRat` model in `RevBayes` (see Data Dryad repository XXXXX and GitHub repository https://github.com/mikeryanmay/musscrat_supp_archive/releases/tag/bioRxiv1.0).

## Statistical Behavior

The `MuSSCRat` model has many parameters relative to the number of observations, 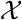 and 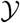. It is therefore unclear how well this complex model can detect rate variation, or distinguish between state-dependent and background sources of rate variation. Accordingly, we performed a simulation study to characterize the statistical behavior of the state-depended model. Specifically, we performed experiments to understand: (1) its ability to detect state-dependent rate variation in the absence of background-rate variation; (2) its ability to detect state-dependent rate variation in the presence of background-rate variation; (3) the cost of including background-rate variation in the model when background rates are actually constant, and; (4) the consequences of assuming background rates are constant when they are actually variable.

For the following analyses, we approximated the joint posterior probability density by running two replicate MCMC simulations for each simulated dataset. We performed MCMC diagnosis to ensure that the joint posterior density was adequately approximated. We provide details of the MCMC simulations and MCMC diagnoses in the Supplemental Material.

### Measures of Performance

#### Frequentist properties

The frequentist interpretation of a Bayesian credible interval (CI) is that the true value of a parameter has a 100(1 − *α*)% chance of being within the 100(1 − *α*)% CI of its corresponding marginal posterior distribution (assuming the model is true, see Huelsenbeck and Rannala 2004). With this interpretation in mind, we assessed the frequentist properties of the 95% CI inferred for our simulated datasets (assuming the conventional significance level, *α* = 0.05). We define the *coverage probability* as the frequency with which the true value of a parameter is contained in the 95% CI, the *false-positive rate* as the frequency with which the true value is excluded from the 95% CI (one minus the coverage probability), and the *power* as the frequency with which a state-independent model is excluded from the 95% CI when the state-dependent model is true.

#### Accuracy and bias

We assess the accuracy and bias of the posterior-mean estimate of 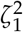 using the percent-error statistic, defined as:

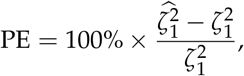

where 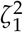 is the true value of state-dependent rate for discrete state 1, and 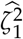 is estimated value of the state-dependent rate for discrete state 1 (we use the mean of the corresponding marginal posterior distribution). Values of PE < 0 indicate an underestimate; conversely, values of PE > 0 indicate an overestimate.

### Simulation Experiments

#### Experiment 1: Constant background rates

We simulated datasets of different sizes (with *c* = {1, 2, 4, 8} continuous characters), over a variety of tree sizes (with *N* = {25, 50, 100} species), and state-dependent rate-ratios, 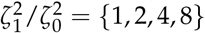 (where 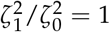 corresponds to the case when rates do not depend on the state of the discrete trait). For each combination of *c*, *N*, and 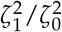, we simulated 100 trees under a constant-rate birth–death process with a speciation rate of 1 and an extinction rate of 0.5 using the `R` (R Core Team 2017) package `TESS` (Höhna 2013; Höhna et al. 2015); we then rescaled each tree to have a root height of 1.

For each tree, Ψ^sim^, we simulated a discrete-character history under a symmetric continuous-time Markov model, with an expectation of five discrete-trait changes. We accomplished this by computing the average tree length for the set of trees simulated under each value of *N*, 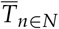, and rescaled the matrix *Q* so that the mean number of changes across trees of size *n* was 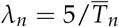. We sampled the discrete state at the root from the stationary distribution of the continuous-time Markov model, and simulated the discrete character forward in time over the phylogeny using the `sim.history()` function in the `R` package `phytools` (Revell 2012b). We conditioned the simulated history on having at least 20% of the tips in each of the discrete states using rejection sampling. We recorded the discrete characters at the tips, 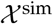, and the discrete-character history, ***κ***^sim^.

For each tree, we used custom `R` scripts to simulate correlation matrices from an LKJ distribution with *η* = 1. For each simulated matrix, we specified the relative rate for each of the *c* continuous characters by sampling each rate from a Gamma(*α* = 4, *β* = 1) distribution, then rescaled the rates to have a mean of 1; this is equivalent to sampling 1/*c*^th^ of the relative rates from a symmetric Dirichlet 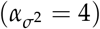 distribution. This simulation scenario explores our ability to detect state-dependent rate variation in the absence of background-rate variation. Accordingly, we specified a background rate of 1 for all lineages; ***β*^2^ = 1**.

For each tree, we used ***κ***^sim^, *R*, and ***σ*^2^**, and the value of 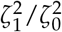 associated with that tree to compute the variance-covariance matrix for each lineage, ∑_*l*_. We then used a custom `R` script to simulate *c* continuous characters, 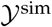, uder the multivariate Brownian motion using the `R` package `mvtnorm` (Genz et al. 2017), assuming that the continuous states at the root of the tree were all 0, ***y***_1_ = **0**.

We analyzed each simulated dataset in `RevBayes` under the `MuSSCRat` model, assuming the true phylogeny was known. We constrained the model so that the background rate was equal for each lineage (*i.e.*, 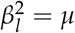 for all *l*), and excluded the standard-deviation parameter, *ν*, from the model. We estimated the remaining model parameters from the simulated data using the priors described in the Bayesian inference section, above. Since the generating model and the inference model both exclude background-rate variation, this simulation scenario reflects the performance of the method when the model is correctly specified.

The false-positive rate for this simulation experiment was 5.3% (*i.e.*, when 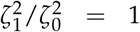; Fig. 4, the top row of the top three panels), which is indistinguishable from the expected 5% (two-tailed binomial test p ≠ 0.05, *p*-value ≈ 0.6). Next, we computed the power when rates of continuous-character evolution varied among discrete states 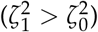. The power ranged from ≈ 21% to 100% from the worst-case to the best-case scenarios. Predictably, power improved as the number of continuous characters and species increased; overall, the average power was 80.4% (Fig. 4, top three panels, excluding the top row). The posterior-mean estimate of 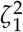 was slightly biased for small numbers of continuous characters (*c* ≤ 2), but quickly converged to the true value as the number of characters and species increased (Fig. 5, top row of panels).

**Figure 4:**
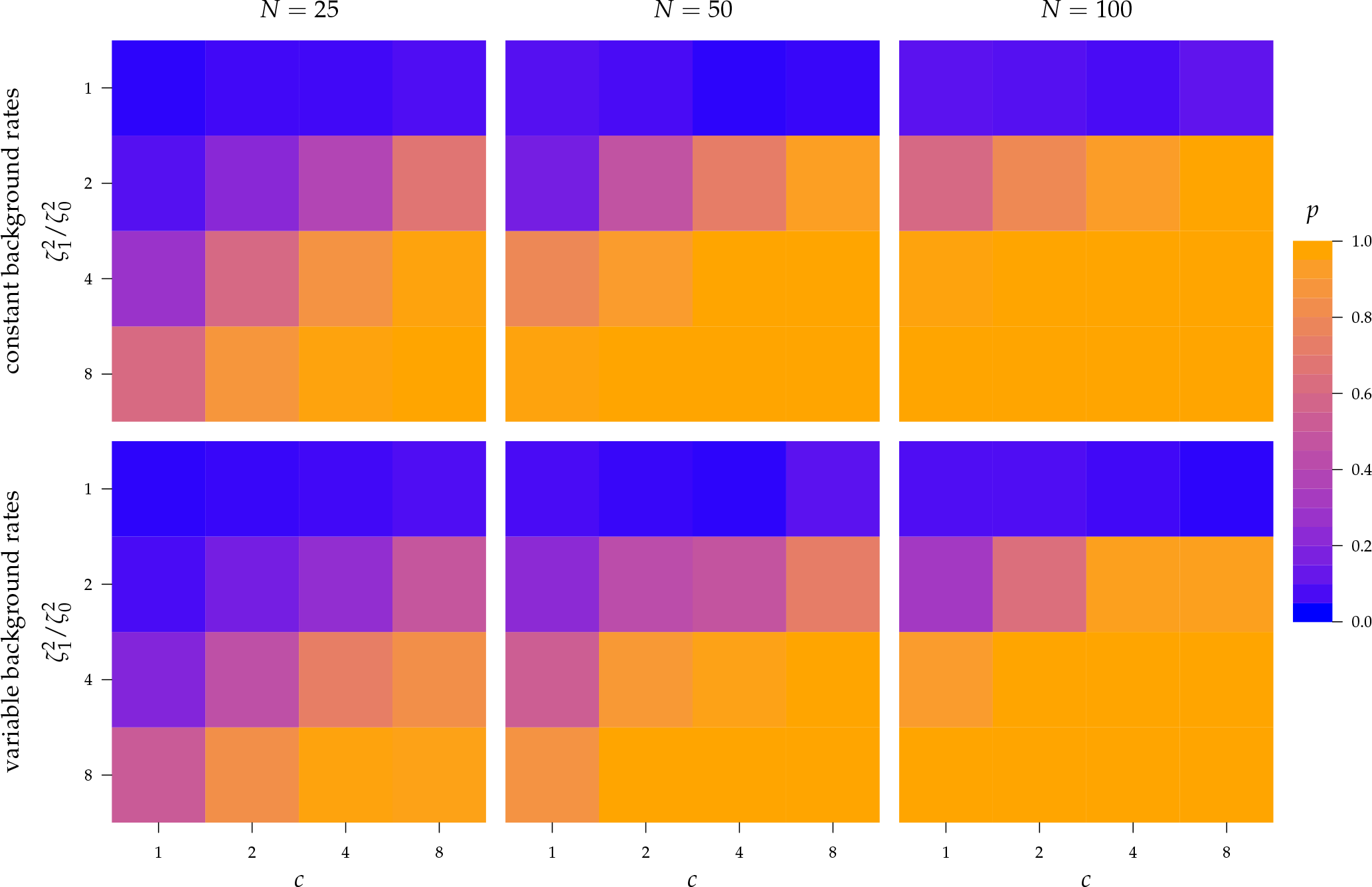
The frequency with which 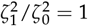 was contained in the 95% credible interval when background rates were constant (top row of panels) or variable (bottom row of panels). Each panel corresponds to simulations for a given number of species, *N*. Within each panel, rows correspond to different degrees of state-dependent rate variation, 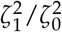, and columns correspond to different numbers of continuous characters, *c*. Each cell presents the fraction of the 95% credible intervals that exclude 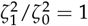, colored according to the scale (at right).

**Figure 5:**
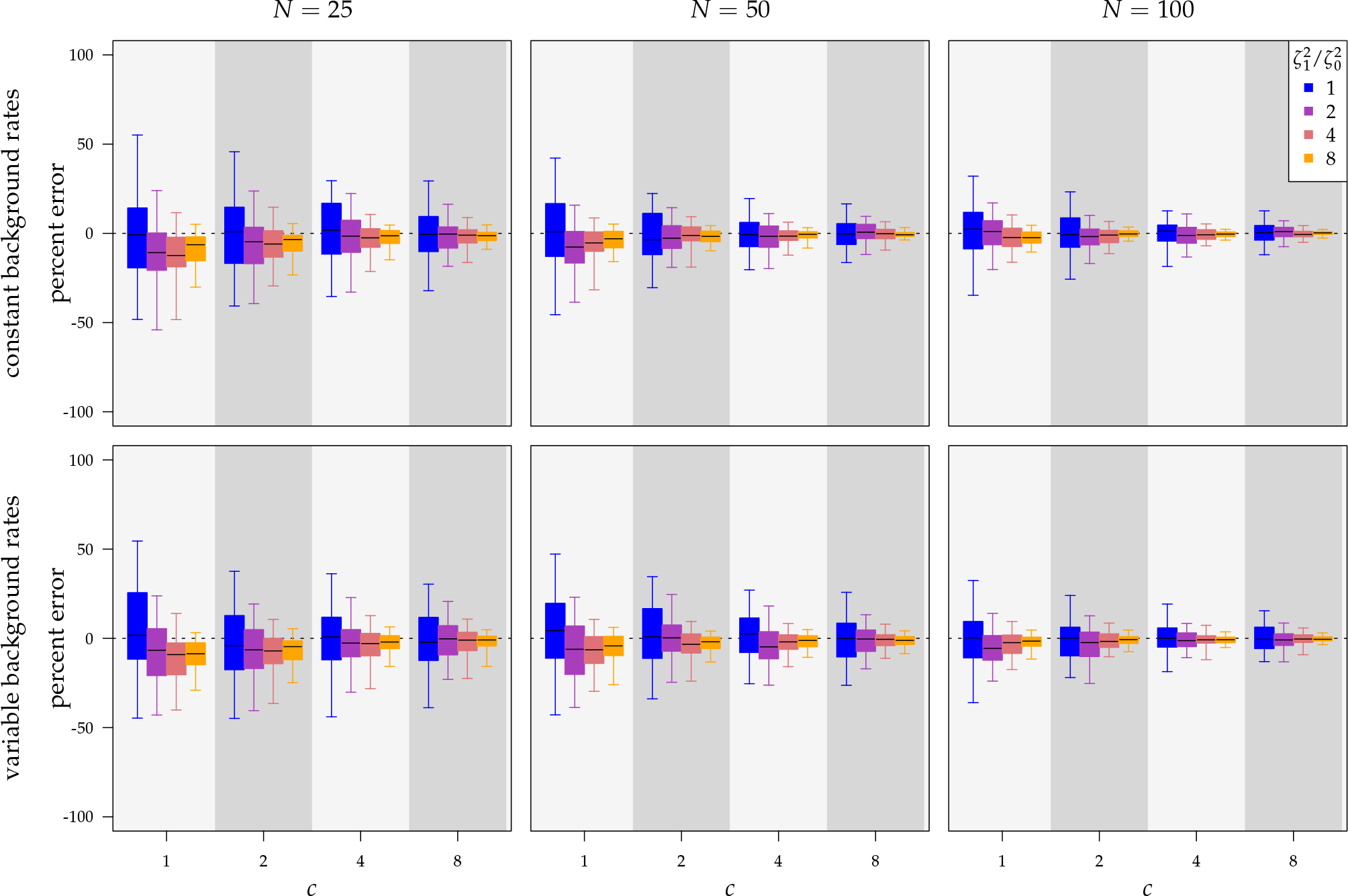
The percent error of the posterior mean estimates of 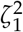 when background rates were constant (top row of panels) or variable (bottom row of panels). Each column of panels corresponds to simulations for a given number of species, *N*. Within each panel, boxplots depict the distribution of percent error across 100 simulated datasets for each of the *c* continuous characters (along the x-axis), colored by the true state-dependent rates, 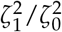 (see inset legend). Boxplots represent the middle 50% (boxes) and the middle 95% (whiskers) of simulations.

#### Experiment 2: Variable background rates

In this simulation scenario, we reused all of the simulated trees, discrete-character histories, correlation matrices, and relative-rate parameters describing the degree of variation among continuous characters from the first simulation experiment (with constant background rates). In this simulation, however, we included variation in the background rates of continuous-character evolution.

For each tree, we simulated lineage-specific rates of continuous-character evolution by drawing the background rate for each lineage, 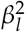, from a lognormal distribution with mean *μ* = 1 and standard deviation *ν* = *H*. We then computed lineage-specific variance-covariance matrices using 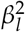 for each lineage, Σ*_l_*, and simulated continuous characters as described in the first simulation experiment.

For this scenario, we analyzed each simulated dataset using the `MuSSCRat` model with background-rate variation, by allowing 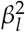 to vary among branches, as described in the Bayesian inference section (`MuSSCRat` + UCLN). Again, this simulation scenario reflects the performance of the method when the model is correctly specified, since the data-generating model and the inference model both allow background rates to vary among lineages.

The false-positive rate for this experiment was 4.9% (Fig. 4, top row of bottom three panels), again indistinguishable from the expected 5% (two-tailed binomial test p ≠ 0.05, *p*-value ≈ 1). The power was only slightly lower than that of experiment 1: the power was 14.7% in the worst case, and 100% in the best case. On average, the power was 73.7% (Fig. 4, bottom three panels, excluding top row). Again, the posterior-mean estimate of 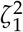 was only modestly biased for analyses based on a small number of continuous characters (Fig. 5, bottom row of panels).

#### Experiment 3: Cost of background-rate variation

When background rates of continuous-character evolution are constant, we expect that including unnecessary parameters (*i.e.*, to accommodate background-rate variation) in the inference model should decrease our ability to detect state-dependent rate variation. The goal of this simulation experiment is to understand the cost of accommodating background-rate variation when it is absent. To achieve this, we reused the datasets from Experiment 1 (simulated under constant background rates) with *N* = 50, *c* = 8, and 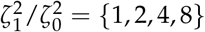, but analyzed these datasets under the `MuSSCRat` + UCLN model.

For Experiment 1—where we correctly assumed that background rates are constant—the coverage probability was 95.3% (two-tailed binomial test p ≠ 0.95, *p*-value ≈ 0.4). By contrast, in this experiment—when we incorrectly assumed that background rates are variable—the coverage probability was 97.4% (two-tailed binomial test p ≠ 0.95, *p*-value = 0.02). Overall, the cost of accommodating background-rate variation when absent was therefore quite modest (97.4% − 95.3% = 2.1%).

#### Experiment 4: Consequences of ignoring background-rate variation

When background rates of continuous-character evolution vary among lineages, we expect that *excluding* background-rate variation from the inference model may be positively misleading. The goal of this simulation experiment is to understand the consequences of failing to accommodate background-rate variation on inferences about state-dependent rates of continuous-character evolution. To achieve this, we reused the datasets from Experiment 2 (simulated under variable background rates), with *N* = 50, *c* = 8, and 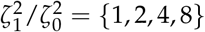, but analyzed these datasets using the “constrained” `MuSSCRat` (*i.e.*, that assumes a constant background rate of evolution by forcing 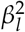 to be the same for all lineages).

For Experiment 2—where we correctly assumed that background rates are variable—the coverage probability was 94.5% (two-tailed binomial test p ≠ 0.95, *p*-value ≈ 0.19). By contrast, in this experiment—when we incorrectly assumed that background rates are constant—the coverage probability decreased to 85.4% (two-tailed binomial test p ≠ 0.95, *p*-value < 1*e* − 10). This decreased coverage probability implies that we are very confident in the wrong answer about 10% more often than we should be. For example, when state-dependent rates are truly equal 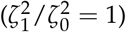, we will incorrectly—but confidently—infer that state-dependent rates differ 17.7% of the time.

## Empirical Analyses

Haemulids (grunts) are a group of percomorph fishes that have previously been used to explore state-dependent rates of continuous-character evolution (Price et al. 2013). Specifically, the hypothesis posits that—owing to the increased habitat complexity of reefs—the feeding apparatus (comprising several continuous traits) of reef-dwelling grunt species should evolve at a higher rate than that of their non-reef-dwelling relatives. We revisit this hypothesis by analyzing the haemulid data from Price et al. (2013) under the `MuSSCRat` model, using a phylogeny estimated from the more extensive molecular dataset from Tavera et al. (2018).

### Phylogenetic Analyses

We assembled a molecular dataset by subsampling the alignments from Tavera et al. (2018) to include only the 49 species represented in our morphological dataset. We estimated a chronogram under a partitioned substitution model assuming an uncorrelated lognormal branch-rate prior model and a sampled birth–death node-age prior model. We performed posterior-predictive tests to ensure that the substitution model provided an adequate description of the substitution process. We computed the maximum *a posteriori* (MAP) chronogram from the posterior distribution of sampled trees and conditioned on this tree in our comparative analyses. We provide details of these analyses in the Supplemental Material.

### Comparative Analyses

We analyzed the continuous morphological data under the `MuSSCRat` model, with habitat type (reef/non-reef) as the discrete character. In these analyses, we conditioned on the MAP chronogram estimated above. We performed a series of analyses to understand: (1) the impact of including or excluding background-rate variation, and; (2) the sensitivity of posterior estimates to the specified priors. For the following analyses, we approximated the joint posterior density by running four replicate MCMC simulations for each analysis using `RevBayes`. Again, we provide details of the MCMC simulations and MCMC diagnoses in the Supplemental Material.

#### Character data

We used eight continuous morphological characters related to the feeding apparatus from Price et al. (2013); we included species that also had molecular sequence data from Tavera et al. (2018), resulting in a total of 49 species. The continuous characters include: (1) the mass of the adductor mandibulae muscle; (2) the length of the ascending process of the premaxilla; (3) the length of the longest gill raker; (4) the diameter of the eye; (5) the length of the buccal cavity; (6) the width of the buccal cavity; (7) the height of the head, and; (8) the length of the head. Rather than size-correcting these characters, we included body size as an additional character (for a total of nine continuous characters). Following Price et al. (2013), we log-transformed each character before the analyses (and cube-rooted the adductor mass prior to log transformation). We used the habitat data from Price et al. (2013) to score each species for the binary discrete character; we coded non-reef-dwelling species and reef-dwelling species as states 0 and 1, respectively.

#### Inferring state-dependent rates

To understand the impact of background-rate variation, we estimated the posterior distribution of the `MuSSCRat` model parameters with and without background-rate variation using the prior settings described in the Bayesian inference section.

The treatment of background-rate variation had a profound impact on both the habitat-specific rate of continuous-character evolution, and also on the inferred history of habitat evolution (Fig. 6). Under the `MuSSCRat` model without background-rate variation, we inferred that the feeding apparatus of reef-dwelling haemulids evolved ≈ 16 times faster than that of their non-reef-dwelling relatives; under the `MuSSCRat` + UCLN model, we inferred a ≈ 2.6-fold increase in the evolutionary rate of reef-dwelling species (95% CIs [10.3 − 23.9] and [1.1 − 5.1], respectively).

**Figure 6:**
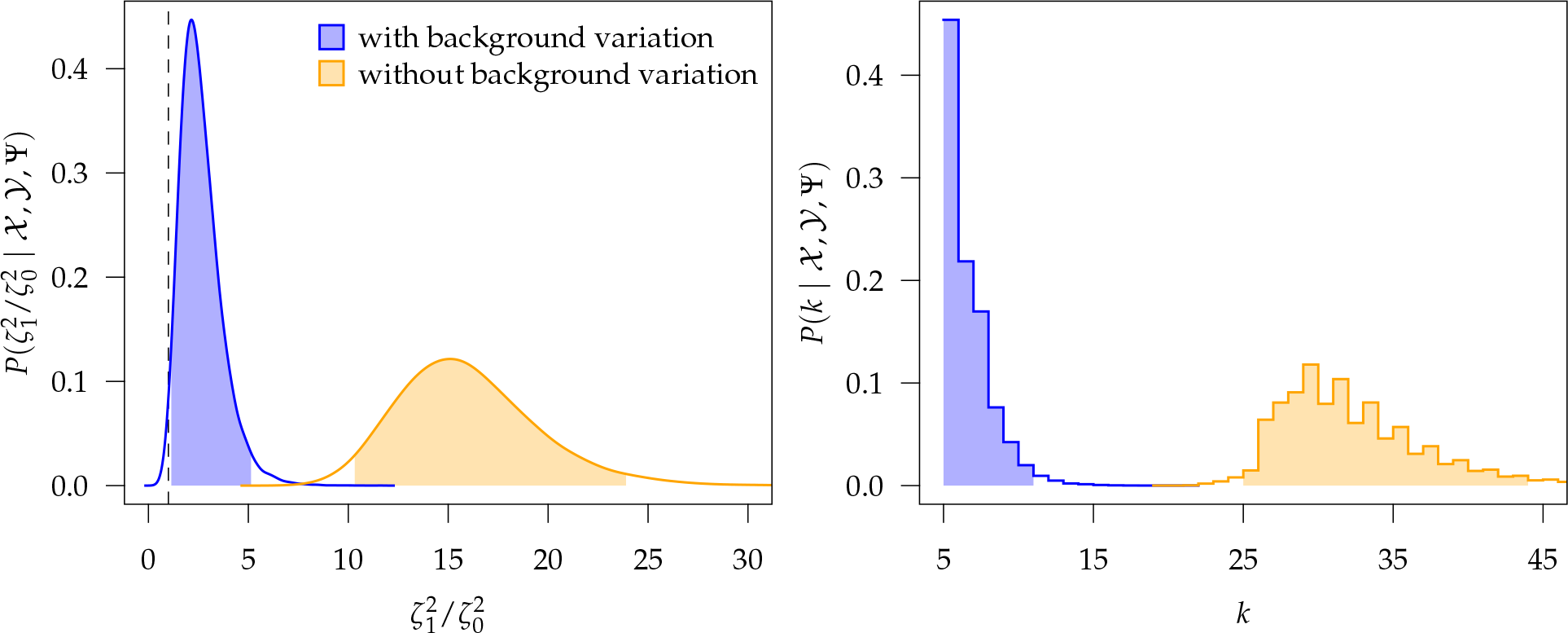
At left, the posterior densities (curves) and the 95% CI (shaded regions) for the state-dependent rate-ratio, 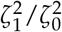, when the background-rates are constant (orange), or when they vary among lineages (blue), inferred for the haemulid dataset. The dashed vertical line corresponds to 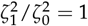. At right, the posterior distribution (lines) and the 95% CI (shaded regions) for the number of habitat transitions, *k*, assuming the background-rates are constant (orange), or vary among lineages (blue).

Examining the posterior distribution of habitat transitions reveals that excluding background-rate variation implies biologically implausible scenarios of habitat evolution. When we disallowed background-rate variation, we inferred ≈ 32 transitions between reef- and non-reef habitats across the phylogeny; when we allowed background rates to vary, we inferred a more reasonable ≈ 6.2 transitions (95% CIs [25 – 43] and [5 – 10], respectively).

#### Prior sensitivity

We assessed the prior sensitivity of inferences by performing a series of analyses using different prior values for various parameters of the model. Specifically, we explored the following prior values:

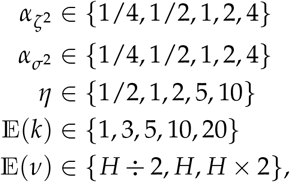

where 𝔼(*ν*) is the prior expected standard deviation of the background-rate variation model. We varied a single prior setting at a time, rather than testing all possible combinations of these priors; we left the remaining priors as described in the Bayesian inference section, for a total of 23 prior combinations.

Most prior settings appear to have little impact on the posterior distribution of the focal parameter, 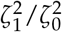 (Fig. 7). Unsurprisingly, the prior on the focal parameter, 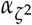, had the greatest influence on the state-dependent rate estimates: the posterior-mean estimate ranged from 2.23 to 2.75 over the priors that we tested (Fig. 7, left band); in all cases 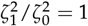 was excluded form the 95% CI. We discuss the (negligible) prior sensitivity of the remaining model parameters in the Supplemental Material.

**Figure 7:**
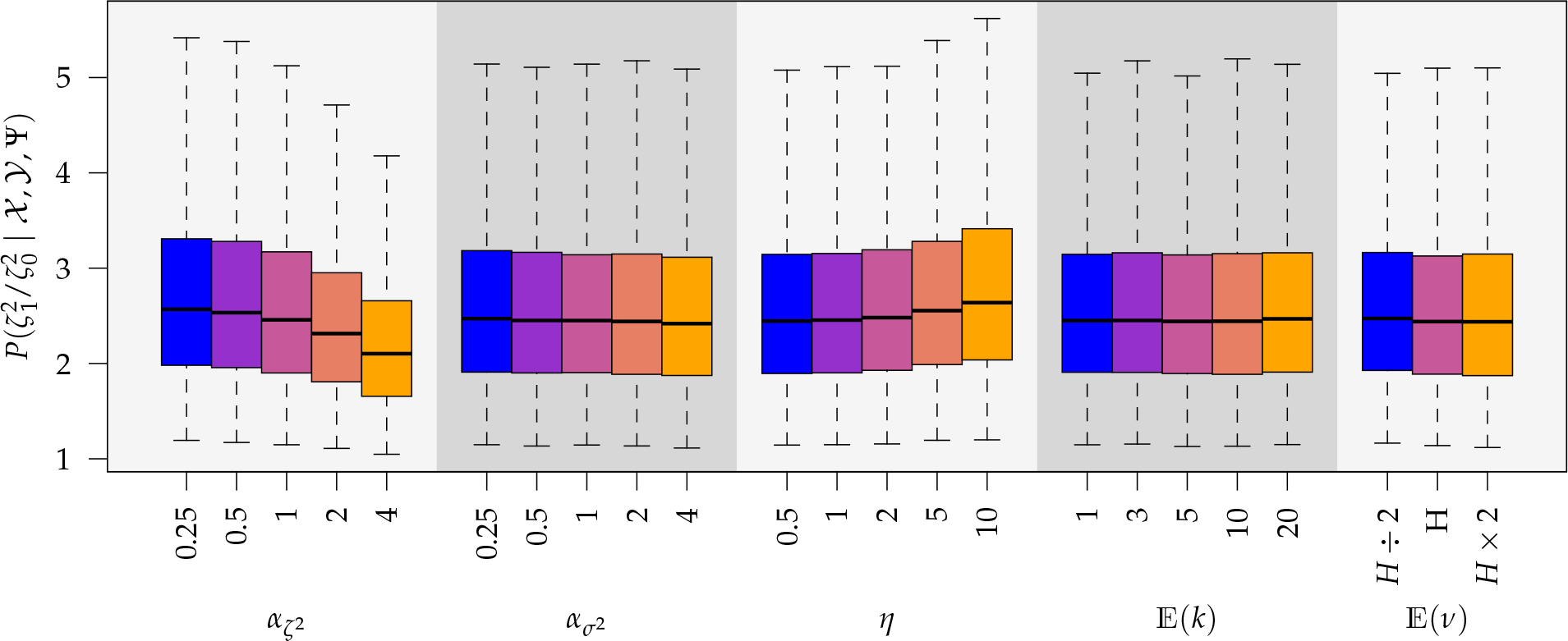
The posterior densities of the state-dependent rate-ratio, 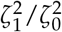, for the haemulid dataset under various priors. Each band of boxplots corresponds to a different prior-sensitivity experiment. Within each band, boxplots represent the 50% CI (box) and 95% CI (whiskers) for the posterior density under a particular value of that prior.

## Discussion

Understanding the factors that drive variation in rates of character evolution is a fundamental goal for evolutionary comparative biologists. Current approaches for assessing the influence of a discrete character on rates of continuous-character evolution suffer from two problems: (1) they do not correctly characterize the mutually informative relationship between the discrete and continuous characters, and (2) they compare against a simple—and likely unrealistic—null model, potentially misleading inferences about state-dependent rates due to rate variation that is unrelated to the discrete character of interest, which we term “background-rate variation”. This second problem is especially concerning, given that rates of evolution are likely to vary greatly across the Tree of Life, and for many reasons not related to the discrete character a particular researcher is investigating.

We present a Bayesian method that deals with both of these issues using a model (`MuSSCRat`) that correctly integrates over discrete character histories with extensions that accommodate background-rate variation. This method involves estimating a large number of parameters, especially compared to the size of typical morphological datasets. This raises serious questions about the reliability of inferences made using the method—especially because the background-rate variation model may wash out any signal of state-dependent rate variation—and also about the sensitivity of inferences to the choice of priors. In the following sections, we describe simulation and empirical results that shed light on the statistical behavior of the method.

### Statistical behavior under simulation

We explored the ability of the `MuSSCRat` model to infer state-dependent rates of continuous-character evolution using simulated data. We varied the simulations over the number of species, the number of continuous characters, and the degree of state-dependent rate variation. We repeated the simulations under different background-rate models: “background-constant” simulations, where background-rates were the same across lineages, and “background-variable” simulations, where background-rates were allowed to vary among lineages.

When the model was correctly specified (*i.e.*, when we inferred parameters under the true background-rate model), the method had appropriate frequentist behavior: the false-positive rate was approximately 5%, and the power increased with the number of taxa, the number of continuous characters, and the degree of state-dependent rate variation. The power was modestly reduced for background-variable simulations (≈ 74%) compared to the background-constant simulations (≈ 80%). Posterior-mean estimates of the state-dependent rate parameters were biased only for small trees (*N* = 25) or datasets with only one or two continuous characters. These results suggest that researchers should be able to reliably infer the state-dependent rate parameters for datasets with a reasonable number of species and continuous characters.

We also used our simulated datasets to assess the costs of including background-rate variation, as well as the consequences of ignoring it. Including unnecessary parameters in the model (overspecification) should lead to increased uncertainty and a concomitant decrease in power: *i.e.*, for background-constant data, allowing for background-rate variation in the inference model should dampen the signal of state dependence. Conversely, excluding parameters from the model (underspecification) should lead to artifactually increased confidence and a higher false-positive rate: *i.e.*, for background-variable data, an inference model that assumes that background rates are constant may spuriously interpret the unmodeled rate variation as additional evidence for state dependence. In our simulations, the cost of model overspecification (an ≈ 2% decrease in power) was minor compared to the consequences of model underspecification (an ≈ 10% increase in the false-positive rate).

### Empirical impact of background-rate variation

We reanalyzed the trophic character data for the haemulids (grunts) from Price et al. (2013) with constant and variable background rates. The inclusion (or exclusion) of background-rate variation in the inference model had a profound impact on inferences regarding the degree of state-dependent rate variation. Under the constant-background-rate model, reef-dwelling species were inferred to evolve more than 15 times faster than their non-reef-dwelling relatives. By contrast, we inferred a ≈ 2.6-fold increase when we allowed background-rate variation. Furthermore, the history of the discrete character inferred under the constant-background-rate model involved an implausible number of habitat transitions. Together, these results suggest that, while trophic-character evolution within the haemulids is elevated within reefs, other factors (manifest as background-rate variation) also played an important role in the evolution of continuous traits in this group.

### Benefits of being Bayesian

Our Bayesian implementation comes with all of the usual advantages of Bayesian inference: the marginal posterior distributions for parameters have a natural interpretation (the 95% credible interval contains the true value of the parameter 95% of the time), and these estimates are automatically averaged over uncertainty in all of the parameters. Indeed, our implementation also allows us to accommodate uncertainty in the tree topology and branch lengths/divergence times, although the impact of phylogenetic uncertainty appears to be relatively mild for haemulids (see Supplemental Material).

Of course, the need to specify prior densities for each model parameter means that posterior estimates may be sensitive to arbitrary prior choices. However, at least for haemulids, inferences about state-dependent rates appear to be quite robust over a range of reasonable priors. We note that our results regarding the impact of phylogenetic uncertainty and prior sensitivity are dataset specific, and may not be generally true for all/most datasets. For this reason, we urge users to perform similar sensitivity assessments for their empirical studies.

Beyond the prosaic strengths and weaknesses of Bayesian inference, adopting a Bayesian framework allowed us to overcome two critical issues for the `MuSSCRat` model. First, the joint distribution of the discrete and continuous characters implied by the `MuSSCRat` model makes it difficult (perhaps impossible) to calculate the full likelihood analytically. Specifying our model in a Bayesian framework allowed us to use a Markov chain Monte Carlo technique—data augmentation—to simplify these likelihood calculations and correctly describe the joint evolution of discrete and continuous characters. We note that similar Monte Carlo integration techniques might be practical for maximum-likelihood applications; indeed, these Monte Carlo solutions have been used to perform maximum-likelihood inference for similar types of problems (Mayrose and Otto 2010; Levy Karin et al. 2017). We conducted experiments that suggest that such approaches would be unreliable for the haemulid dataset (see Supplemental Material), but more work is necessary to understand the generality of these results.

Second, adopting a Bayesian framework allowed us to include background-rate variation while retaining the ability to perform reliable inference under the model. The `MuSSCRat` model with background-rate variation is inherently nonidentifiable: multiple combinations of background-rate and state-dependent-rate parameters can have identical likelihoods, so a unique maximum-likelihood estimate of the parameters may not exist. Within a Bayesian setting, the joint prior distribution acts to “tease apart” combinations of parameters that would otherwise have identical likelihoods, thus making it possible to infer parameters under nonidentifiable models.

### Broader context

There is mounting concern within the phylogenetic comparative community that methods for understanding relationships between evolutionary variables are unreliable (Beaulieu et al. 2013; Maddison and FitzJohn 2014; Rabosky and Goldberg 2015; Beaulieu and O’Meara 2016; Uyeda et al. 2018). In particular, Uyeda et al. (2018) and Beaulieu and O’Meara (2016) identify two (not mutually exclusive) concerns, namely: (1) our models fail to account for “unreplicated evolutionary events”; and (2) our model-selection procedures often involve comparisons against an extremely unrealistic—and therefore easy-to-reject—null model.

We agree with Uyeda et al. (2018) that unmodeled rare events can be problematic; however, it is clear that unmodeled *common* events may be similarly treacherous. Our simulation study included datasets where background rates of continuous-trait evolution varied continuously across branches of the tree, *i.e.*, background-rate variation was pervasive. We demonstrated that ignoring this pervasive background-rate variation compromised our ability to accurately infer the rate of state-dependent evolution: in ≈ 15% of these analyses, the true value of state-dependent rate variation was excluded from the 95% CI. We believe that it is important to accommodate “nuisance” rate variation, whether it is rare or common.

Beaulieu and O’Meara (2016) clarify the fundamental issue raised by Rabosky and Goldberg (2015): when asked to choose between two models, it should come as no surprise when model-comparison procedures reject an overly simplistic, constant-rate null model in favor of a very specific, variable-rate alternative model. The danger is that *any* rate variation (whether or not it is associated with a focal variable of interest) will be interpreted by a model-comparison procedure as evidence against a constant-rate null model. This logic also applies to parameter-estimation procedures: when considering a variable-rate model with a single explanatory variable, it seems likely that any evidence for heterogeneity is at risk of being spuriously attributed to the factor of interest. We refer to this problem as the “straw-man effect”, and we suspect that it applies whenever model comparison includes at least one constant-rate model or—in the case of parameter estimation—whenever a variable-rate model is overly simplistic.

A possible solution to the straw-man effect—one that we favor, but by no means the only conceivable solution—is to move away from hypothesis testing and overly specific variable-rate models toward explicit models of background variation, as in Beaulieu and O’Meara (2016) and the present work. A justifiable concern with such models is that the results may be sensitive to the assumed nature of the background variation (whether it is due to hidden characters, lineage-specific effects, the environment, or any number of alternatives), which may be difficult to justify *a priori* or distinguish *a posteriori*.

### Future prospects

Our `MuSSCRat` model makes many simplifying assumptions about the nature of the evolutionary process under consideration. For example, we assume that the continuous characters evolve under a Brownian motion process; that the underlying evolutionary variance-covariance matrix, Σ, is the same over the entire tree; that the discrete characters evolve under a simple continuous-time Markov process; and that the background-rate variation is adequately described by an uncorrelated lognormal distribution. Each of these assumptions provides opportunities for future model development.

There are many alternatives to the Brownian motion model in the phylogenetic comparative toolkit, perhaps chief among them the Ornstein-Uhlenbeck (OU) process (Hansen 1997; Butler and King 2004). State-dependent OU process models have been widely used to detect shifts in evolutionary optima associated with discrete characters. It seems likely that these approaches are vulnerable to unmodeled process heterogeneity, since they typically compare state-dependent models against homogeneous models (the “straw-man effect”). Extending the basic framework presented here to the OU process should be straightforward, and may help ameliorate the issues raised by Uyeda et al. (2018).

We have assumed that the “structure” of the evolutionary variance-covariance matrix is independent of the discrete character (in the sense that the discrete character affects only the magnitude of the variance-covariance matrix). Recently, Caetano and Harmon (2018) developed a method that allows the structure of the variance-covariance matrix to depend on the state of a discrete character. While this is an important advance over models that assume that the variance-covariance matrix is homogeneous, it too may suffer from the straw-man effect: heterogeneity in the variance-covariance matrix unrelated to the discrete character may be positively misleading. Extending the framework developed by Caetano and Harmon (2018) to incorporate background variation in the variance-covariance matrix is an important avenue for future development.

We have employed a relatively simple continuous-time Markov process to model the evolution of the discrete character. This is of potential concern for haemulids, where there is strong evidence that rates of lineage diversification (speciation – extinction) are elevated in reef-dwelling species (data not shown). The extent to which state-dependent lineage diversification models impact the distribution of discrete-character histories—and therefore compromise inferences about discrete-state-dependent rates of continuous-character evolution—is an open question. Although the modeling task is relatively straightforward (FitzJohn 2010), developing the computational machinery to do inference under such models—where a discrete character affects both rates of continuous-character evolution *and* rates of lineage diversification—is a nontrivial technical challenge.

Finally, we have assumed that background-rate variation follows an uncorrelated lognormal prior distribution. Because the `MuSSCRat` model with background-rate variation is nonidentifiable, specifying a prior on background-rate variation is critical to teasing apart the relative contributions of background- and state-dependent effects to the overall rate variation. However, the Bayesian solution to nonidentifiability is not without caveats: since we rely on the prior to distinguish the relative effects of background-rate variation and state-dependent rates, our inferences about state-dependent rates may be sensitive to the assumed model of background-rate variation. Unfortunately, it is not clear that alternative models of background-rate variation can be reliably distinguished using standard model-selection procedures. However, Bayesian models provide a natural way for updating our prior beliefs and integrating information across disparate datasets, and we foresee a fruitful Bayesian research program that leverages information from across the Tree of Life (extinct and extant) to characterize overall heterogeneity in the evolutionary process, and to more accurately distinguish among its multifarious causes.

## Supplementary Material

Supplementary scripts and data (including the Haemulidae data and simulated data used in this study) can be found in the Data Dryad repository DOI:X and the GitHub repository https://github.com/mikeryanmay/musscrat_supp_archive/releases/tag/bioRxiv1.0.

## Acknowledgements

We would like to thank Bruce Rannala, Jiansi Gao, Nikolai Vetr, Sebastian Höhna, Michael Landis, Peter Wain-wright and Samantha Price for their thoughtful and generous discussion throughout the process of writing this manuscript.

