## supplemental material for "A Bayesian Approach for Inferring the Impact of a Discrete Character on Rates of Continuous-Character Evolution in the Presence of Background-Rate Variation"

### Supplemental Material for: Bayesian Estimation of Discrete-State-Dependent Rates of Continuous-Character Evolution

#### Contents

**S.1 Derivation of the Transition Probability Density** **S2**
S.1.1 Finite Difference Equations . . . S3
S.1.2 Taylor Series Expansion . . . S3
S.1.3 Limit of the Finite Difference Equations . . . S4
S.1.4 Partial Differential Equation for the Transition Probability Density . . . S8

**S.2 Identifiability of the State-Dependent Model** **S9**

**S.3 Numerical Consequences of Sequential Analysis** **S10**

**S.4 Standard Deviation of the Lognormal Distribution** **S14**

**S.5 MCMC Proposals** **S15**
S.5.1 Reversible-jump proposals for symmetric and asymmetric models . . . S15
S.5.2 Proposals on discrete-character histories . . . S15

**S.6 Empirical Analyses** **S17**
S.6.1 MCMC Diagnosis . . . S17
S.6.2 Model Adequacy . . . S17
S.6.3 Phylogenetic Analyses . . . S17
S.6.4 Joint Analysis of Phylogeny and Character Evolution . . . S22
S.6.5 Prior Sensitivity . . . S23

**S.7 Simulation Study** **S28**
S.7.1 MCMC Analyses and Diagnosis . . . S28
S.7.2 Statistical Behavior . . . S28

#### S.1 Derivation of the Transition Probability Density

We derive a set of partial differential equations that describe the evolution of a set of characters under the state-dependent multivariate Brownian motion model (the Kolmogorov backward equations); these equations are a special case of equations (B.9b) from FitzJohn (2010), but we provide the derivation here for the sake of completeness and notational consistency.

We assume that there are $c$ continuous characters (each of which may be any real number) and a single discrete character (with $d$ states). We denote the joint probability density of the continuous characters, $\boldsymbol{y}$, and the $i^{\text{th}}$ discrete character state at time $t$ as $P_i(\boldsymbol{y}, t)$. To compute the joint probability density at some infinitesimal time later, $P_i(\boldsymbol{y}, t + \Delta t)$, we consider all possible changes in the characters in the interval $\Delta t$ and compute their corresponding probability density (assuming $\Delta t$ is sufficiently small so that the probability of two changes in the discrete character is on the order of $\Delta t^2$ and can safely be ignored). Specifically,

$$\begin{aligned}
P_i(\boldsymbol{y}, t + \Delta t) = &\sum_{j \neq i}^{d} q_{ji} \Delta t \int\limits_{\boldsymbol{z} \in \mathbb{R}^c} f(\boldsymbol{z}, t \mid \boldsymbol{y}, t + \Delta t) P_j(\boldsymbol{z}, t) \mathrm{d}\boldsymbol{z} \\
&- \sum_{j \neq i}^{d} q_{ij} \Delta t \int\limits_{\boldsymbol{z} \in \mathbb{R}^c} f(\boldsymbol{z}, t \mid \boldsymbol{y}, t + \Delta t) P_i(\boldsymbol{z}, t) \mathrm{d}\boldsymbol{z} \\
&+ \int\limits_{\boldsymbol{z} \in \mathbb{R}^c} f(\boldsymbol{z}, t \mid \boldsymbol{y}, t + \Delta t) P_i(\boldsymbol{z}, t) \mathrm{d}\boldsymbol{z} \\
&+ O(\Delta t^2),
\end{aligned}$$

where $f(\cdot)$ is the transition probability density of the multivariate Brownian motion within a given discrete state (*i.e.*, a multivariate normal density with mean $\boldsymbol{y}$ and variance $\Sigma_i \Delta t$). The multidimensional integral,

$$\int\limits_{\boldsymbol{z} \in \mathbb{R}^c} f(\boldsymbol{z}, t \mid \boldsymbol{y}, t + \Delta t) P_i(\boldsymbol{z}, t) \mathrm{d}\boldsymbol{z} = \int_{-\infty}^{\infty} \cdots \int_{-\infty}^{\infty} f(\boldsymbol{z}, t \mid \boldsymbol{y}, t + \Delta t) P_i(\boldsymbol{z}, t) \mathrm{d}z_1 \cdots \mathrm{d}z_c,$$

represents the probability of evolving into state $\boldsymbol{y}$ integrating over all starting states, $\boldsymbol{z}$, in proportion to their initial probability, $P_i(\boldsymbol{z}, t)$.

##### S.1.1 Finite Difference Equations

We derive the system of partial differential equations by taking the limit of the *finite difference equations* as the interval $\Delta t$ approaches zero. The finite difference equation for discrete state $i$ is:

$$\begin{aligned}\frac{P_i(\boldsymbol{y},t+\Delta t)-P_i(\boldsymbol{y},t)}{\Delta t} = &\sum_{j\neq i}^{d} q_{ji} \int\limits_{\boldsymbol{z}\in\mathbb{R}^c} f(\boldsymbol{z},t \mid \boldsymbol{y},t+\Delta t) P_j(\boldsymbol{z},t)\mathrm{d}\boldsymbol{z} \\ &-\sum_{j\neq i}^{d} q_{ij} \int\limits_{\boldsymbol{z}\in\mathbb{R}^c} f(\boldsymbol{z},t \mid \boldsymbol{y},t+\Delta t) P_i(\boldsymbol{z},t)\mathrm{d}\boldsymbol{z} \\ &+\frac{1}{\Delta t} \int\limits_{\boldsymbol{z}\in\mathbb{R}^c} f(\boldsymbol{z},t \mid \boldsymbol{y},t+\Delta t) P_i(\boldsymbol{z},t)\mathrm{d}\boldsymbol{z} \\ &-\frac{1}{\Delta t} P_i(\boldsymbol{y},t) \\ &+O(\Delta t).\end{aligned} \tag{S.1}$$

Before taking the limit of the above difference equations with respect to time, we note that

$$\begin{aligned}P_i(\boldsymbol{y},t) &= \int\limits_{\boldsymbol{z}\in\mathbb{R}^c} f(\boldsymbol{z},t \mid \boldsymbol{y},t+\Delta t) P_i(\boldsymbol{y},t)\mathrm{d}\boldsymbol{z} \\ &= P_i(\boldsymbol{y},t) \int\limits_{\boldsymbol{z}\in\mathbb{R}^c} f(\boldsymbol{z},t \mid \boldsymbol{y},t+\Delta t)\mathrm{d}\boldsymbol{z} \\ &= P_i(\boldsymbol{y},t),\end{aligned} \tag{S.2}$$

since $f(\boldsymbol{z},t \mid \boldsymbol{y},t+\Delta t)$ is a probability density function, and therefore must integrate to 1. Substituting eq. (S.2) into eq. (S.1) allows us to combine the third and fourth terms in eq. (S.1), which yields:

$$\begin{aligned}\frac{P_i(\boldsymbol{y},t+\Delta t)-P_i(\boldsymbol{y},t)}{\Delta t} = &\sum_{j\neq i}^{d} q_{ji} \int\limits_{\boldsymbol{z}\in\mathbb{R}^c} f(\boldsymbol{z},t \mid \boldsymbol{y},t+\Delta t) P_j(\boldsymbol{z},t)\mathrm{d}\boldsymbol{z} \\ &-\sum_{j\neq i}^{d} q_{ij} \int\limits_{\boldsymbol{z}\in\mathbb{R}^c} f(\boldsymbol{z},t \mid \boldsymbol{y},t+\Delta t) P_i(\boldsymbol{z},t)\mathrm{d}\boldsymbol{z} \\ &+\frac{1}{\Delta t} \int\limits_{\boldsymbol{z}\in\mathbb{R}^c} f(\boldsymbol{z},t \mid \boldsymbol{y},t+\Delta t) (P_i(\boldsymbol{z},t)-P_i(\boldsymbol{y},t))\mathrm{d}\boldsymbol{z} \\ &+O(\Delta t).\end{aligned} \tag{S.3}$$

##### S.1.2 Taylor Series Expansion

Next, we take the limit of the finite difference equation as $\Delta t \to 0$. To solve this limit, we expand $P_i(\boldsymbol{z},t)$ as a Taylor series about the point $\boldsymbol{z}=\boldsymbol{y}$. In vector notation, this Taylor expansion (ignoring higher order terms) is:

$$P_i(\boldsymbol{z},t) = P_i(\boldsymbol{y},t) + (\boldsymbol{z}-\boldsymbol{y}) \cdot \nabla P_i(\boldsymbol{y},t) + \frac{1}{2}(\boldsymbol{z}-\boldsymbol{y}) \cdot \boldsymbol{H}_{P_i}(\boldsymbol{y}) \cdot (\boldsymbol{z}-\boldsymbol{y}) + O((\Delta \boldsymbol{y})^3) \tag{S.4}$$

where $\cdot$ is the dot product, $\nabla P_i(\boldsymbol{y}, t)$ is the (vector-valued) gradient (*i.e.*, the multidimensional derivative) of the probability density at $\boldsymbol{y}$:

$$\nabla P_i(\boldsymbol{y}, t) = \left[\frac{\partial P_i}{\partial y_1}, \frac{\partial P_i}{\partial y_2}, \ldots, \frac{\partial P_i}{\partial y_c}\right],$$

$\boldsymbol{H}_{P_i}$ is the Hessian matrix, which contains the second partial derivatives of the probability density:

$$\boldsymbol{H}_{P_i} = \begin{bmatrix} \frac{\partial^2 P_i}{\partial y_1^2} & \frac{\partial^2 P_i}{\partial y_1 \partial y_2} & \cdots & \frac{\partial^2 P_i}{\partial y_1 \partial y_c} \\ \frac{\partial^2 P_i}{\partial y_2 y_1} & \frac{\partial^2 P_i}{\partial y_2^2} & \cdots & \frac{\partial^2 P_i}{\partial y_2 \partial y_c} \\ \vdots & \vdots & \ddots & \vdots \\ \frac{\partial^2 P_i}{\partial y_c \partial y_1} & \frac{\partial^2 P_i}{\partial y_c \partial y_2} & \cdots & \frac{\partial^2 P_i}{\partial y_c^2} \end{bmatrix},$$

$\boldsymbol{H}_{P_i}(\boldsymbol{y})$ is the Hessian matrix evaluated at $\boldsymbol{y}$, and the term $O((\Delta \boldsymbol{y})^3)$ contains terms that involve larger-order changes in $\boldsymbol{y}$ (which we will safely ignore later).

##### S.1.3 Limit of the Finite Difference Equations

The limit of the finite difference equation as $\Delta t \to 0$ is:

$$\begin{aligned} \lim_{\Delta t \to 0} \frac{P_i(\boldsymbol{y}, t + \Delta t) - P_i(\boldsymbol{y}, t)}{\Delta t} = & \lim_{\Delta t \to 0} \sum_{j \neq i}^{d} q_{ji} \int_{\boldsymbol{z} \in \mathbb{R}^c} f(\boldsymbol{z}, t \mid \boldsymbol{y}, t + \Delta t) P_j(\boldsymbol{z}, t) \mathrm{d}\boldsymbol{z} \\ & - \lim_{\Delta t \to 0} \sum_{j \neq i}^{d} q_{ij} \int_{\boldsymbol{z} \in \mathbb{R}^c} f(\boldsymbol{z}, t \mid \boldsymbol{y}, t + \Delta t) P_i(\boldsymbol{z}, t) \mathrm{d}\boldsymbol{z} \\ & + \lim_{\Delta t \to 0} \frac{1}{\Delta t} \int_{\boldsymbol{z} \in \mathbb{R}^c} f(\boldsymbol{z}, t \mid \boldsymbol{y}, t + \Delta t) (P_i(\boldsymbol{z}, t) - P_i(\boldsymbol{y}, t)) \mathrm{d}\boldsymbol{z} \\ & + \lim_{\Delta t \to 0} O(\Delta t). \end{aligned} \tag{S.5}$$

###### S.1.3.1 First term

We solve the terms on the right-hand-side of eq. (S.5) one at a time. Using the Taylor expansion, eq. (S.4), the first term can be re-written:

$$\begin{aligned} & \lim_{\Delta t \to 0} \sum_{j \neq i}^{d} q_{ji} \int_{\boldsymbol{z} \in \mathbb{R}^c} f(\boldsymbol{z}, t \mid \boldsymbol{y}, t + \Delta t) P_j(\boldsymbol{z}, t) \mathrm{d}\boldsymbol{z} \\ & = \lim_{\Delta t \to 0} \sum_{j \neq i}^{d} q_{ji} \int_{\boldsymbol{z} \in \mathbb{R}^c} f(\boldsymbol{z}, t \mid \boldsymbol{y}, t + \Delta t) \Bigg[ P_j(\boldsymbol{y}, t) + (\boldsymbol{z} - \boldsymbol{y}) \cdot \nabla P_j(\boldsymbol{y}, t) \\ & \qquad + \frac{1}{2} (\boldsymbol{z} - \boldsymbol{y}) \cdot \boldsymbol{H}_{P_j}(\boldsymbol{y}) \cdot (\boldsymbol{z} - \boldsymbol{y}) + O((\Delta \boldsymbol{y})^3) \Bigg] \mathrm{d}\boldsymbol{z}. \end{aligned}$$

By expanding the integral, we obtain

$$
\begin{aligned}
\lim_{\Delta t \to 0} \sum_{j \neq i}^{d} q_{ji} \int\limits_{z \in \mathbb{R}^c} & f(\boldsymbol{z}, t \mid \boldsymbol{y}, t + \Delta t) P_j(\boldsymbol{z}, t) \mathrm{d}\boldsymbol{z} \\
&= \sum_{j \neq i}^{d} q_{ji} \Bigg[ \lim_{\Delta t \to 0} \int\limits_{z \in \mathbb{R}^c} f(\boldsymbol{z}, t \mid \boldsymbol{y}, t + \Delta t) P_j(\boldsymbol{y}, t) \mathrm{d}\boldsymbol{z} \\
&\qquad + \lim_{\Delta t \to 0} \int\limits_{z \in \mathbb{R}^c} f(\boldsymbol{z}, t \mid \boldsymbol{y}, t + \Delta t) (\boldsymbol{z} - \boldsymbol{y}) \cdot \nabla P_j(\boldsymbol{y}, t) \mathrm{d}\boldsymbol{z} \\
&\qquad + \lim_{\Delta t \to 0} \int\limits_{z \in \mathbb{R}^c} f(\boldsymbol{z}, t \mid \boldsymbol{y}, t + \Delta t) \frac{1}{2} (\boldsymbol{z} - \boldsymbol{y}) \cdot \boldsymbol{H}_{P_j}(\boldsymbol{y}) \cdot (\boldsymbol{z} - \boldsymbol{y}) \mathrm{d}\boldsymbol{z} \\
&\qquad + \lim_{\Delta t \to 0} O((\Delta \boldsymbol{y})^3) \Bigg].
\end{aligned} \tag{S.6}
$$

The first term on the right-hand-side of eq. (S.6) can be rewritten as:

$$
\begin{aligned}
\lim_{\Delta t \to 0} \int\limits_{z \in \mathbb{R}^c} f(\boldsymbol{z}, t \mid \boldsymbol{y}, t + \Delta t) P_j(\boldsymbol{y}, t) \mathrm{d}\boldsymbol{z} &= P_j(\boldsymbol{y}, t) \lim_{\Delta t \to 0} \int\limits_{z \in \mathbb{R}^c} f(\boldsymbol{z}, t \mid \boldsymbol{y}, t + \Delta t) \mathrm{d}\boldsymbol{z} \\
&= P_j(\boldsymbol{y}, t),
\end{aligned}
$$

since

$$
\int\limits_{z \in \mathbb{R}^c} f(\boldsymbol{z}, t \mid \boldsymbol{y}, t + \Delta t) \mathrm{d}\boldsymbol{z} = 1.
$$

Under multivariate Brownian motion, the probability density of a transition away from $\boldsymbol{y}$ as $\Delta t \to 0$ becomes vanishingly small:

$$
\lim_{\Delta t \to 0} f(\boldsymbol{z}, t \mid \boldsymbol{y}, t + \Delta t) = \delta(\boldsymbol{z} - \boldsymbol{y}),
$$

where $\delta(\cdot)$ represents the Dirac delta function, which assigns infinity probability density to a particular value, $\boldsymbol{y} = \boldsymbol{0}$. This implies that

$$
\int\limits_{z \in \mathbb{R}^c} f(\boldsymbol{z}, t \mid \boldsymbol{y}, t + \Delta t) (\boldsymbol{z} - \boldsymbol{y})^k \mathrm{d}\boldsymbol{z} = 0, \; k > 0.
$$

As a consequence, the final three terms of eq. (S.6) vanish. Ultimately, eq. (S.6) is

$$
\lim_{\Delta t \to 0} \sum_{j \neq i}^{d} q_{ji} \int\limits_{z \in \mathbb{R}^c} f(\boldsymbol{z}, t \mid \boldsymbol{y}, t + \Delta t) P_j(\boldsymbol{z}, t) \mathrm{d}\boldsymbol{z} = \sum_{j \neq i}^{d} q_{ji} P_j(\boldsymbol{y}, t). \tag{S.7}
$$

###### S.1.3.2 Second term

We can use the same process as above to find the limit of the second term of eq. (S.5):

$$
\lim_{\Delta t \to 0} \sum_{j \neq i}^{d} q_{ij} \int\limits_{z \in \mathbb{R}^c} f(\boldsymbol{z}, t \mid \boldsymbol{y}, t + \Delta t) P_j(\boldsymbol{z}, t) \mathrm{d}\boldsymbol{z} = \sum_{j \neq i}^{d} q_{ij} P_i(\boldsymbol{y}, t). \tag{S.8}
$$

###### S.1.3.3 Third term

The final term of eq. (S.5), after substituting the Taylor expansion, becomes:

$$\begin{aligned}
&\lim_{\Delta t \to 0} \frac{1}{\Delta t} \int\limits_{\boldsymbol{z} \in \mathbb{R}^c} f(\boldsymbol{z}, t \mid \boldsymbol{y}, t+\Delta t)(P_i(\boldsymbol{z}, t) - P_i(\boldsymbol{y}, t))\mathrm{d}\boldsymbol{z} \\
&\qquad = \lim_{\Delta t \to 0} \frac{1}{\Delta t} \int\limits_{\boldsymbol{z} \in \mathbb{R}^c} f(\boldsymbol{z}, t \mid \boldsymbol{y}, t+\Delta t)(\boldsymbol{z} - \boldsymbol{y}) \cdot \nabla P_i(\boldsymbol{y}, t)\mathrm{d}\boldsymbol{z} \\
&\qquad + \lim_{\Delta t \to 0} \frac{1}{\Delta t} \int\limits_{\boldsymbol{z} \in \mathbb{R}^c} f(\boldsymbol{z}, t \mid \boldsymbol{y}, t+\Delta t)\frac{1}{2}(\boldsymbol{z} - \boldsymbol{y}) \cdot \boldsymbol{H}_{P_i}(\boldsymbol{y}) \cdot (\boldsymbol{z} - \boldsymbol{y})\mathrm{d}\boldsymbol{z} \\
&\qquad + \lim_{\Delta t \to 0} O((\Delta \boldsymbol{y})^3).
\end{aligned} \tag{S.9}$$

Unpacking the vector notation, the first term of eq. (S.9) can be written:

$$\begin{aligned}
&\lim_{\Delta t \to 0} \frac{1}{\Delta t} \int\limits_{\boldsymbol{z} \in \mathbb{R}^c} f(\boldsymbol{z}, t \mid \boldsymbol{y}, t+\Delta t)(\boldsymbol{z} - \boldsymbol{y}) \cdot \nabla P_i(\boldsymbol{y}, t)\mathrm{d}\boldsymbol{z} \\
&\qquad = \lim_{\Delta t \to 0} \frac{1}{\Delta t} \int\limits_{\boldsymbol{z} \in \mathbb{R}^c} f(\boldsymbol{z}, t \mid \boldsymbol{y}, t+\Delta t)(z_1 - y_1)\frac{\partial P_i}{\partial y_1}\mathrm{d}\boldsymbol{z} \\
&\qquad + \lim_{\Delta t \to 0} \frac{1}{\Delta t} \int\limits_{\boldsymbol{z} \in \mathbb{R}^c} f(\boldsymbol{z}, t \mid \boldsymbol{y}, t+\Delta t)(z_2 - y_2)\frac{\partial P_i}{\partial y_2}\mathrm{d}\boldsymbol{z} \\
&\qquad \vdots \\
&\qquad + \lim_{\Delta t \to 0} \frac{1}{\Delta t} \int\limits_{\boldsymbol{z} \in \mathbb{R}^c} f(\boldsymbol{z}, t \mid \boldsymbol{y}, t+\Delta t)(z_c - y_c)\frac{\partial P_i}{\partial y_c}\mathrm{d}\boldsymbol{z}
\end{aligned} \tag{S.10}$$

The transition density of the $j^{\text{th}}$ character, marginal with respect to the remaining characters, is:

$$f(z_j, t \mid y_j, t+\Delta t) = \int\limits_{\boldsymbol{z}'} f(\boldsymbol{z}, t \mid \boldsymbol{y}, t+\Delta t)\mathrm{d}\boldsymbol{z}', \tag{S.11}$$

with $\boldsymbol{z} = z_j \cup \boldsymbol{z}'$, and the (multidimensional) integral taken over all possible values of $\boldsymbol{z}' \in \mathbb{R}^{c-1}$. We can then write the $j^{\text{th}}$ term on the right-hand-side of eq. (S.10) as:

$$\begin{aligned}
&\lim_{\Delta t \to 0} \frac{1}{\Delta t} \int\limits_{\boldsymbol{z} \in \mathbb{R}^c} f(\boldsymbol{z}, t \mid \boldsymbol{y}, t+\Delta t)(z_j - y_j)\frac{\partial P_i}{\partial y_j}\mathrm{d}\boldsymbol{z} \\
&\qquad = \frac{\partial P_i}{\partial y_j} \lim_{\Delta t \to 0} \frac{1}{\Delta t} \int\limits_{\boldsymbol{y} \in \mathbb{R}^c} (z_j - y_j) f(\boldsymbol{z}, t \mid \boldsymbol{y}, t+\Delta t)\mathrm{d}\boldsymbol{z} \\
&\qquad = \frac{\partial P_i}{\partial y_j} \lim_{\Delta t \to 0} \frac{1}{\Delta t} \int\limits_{y_j} (z_j - y_j) f(z_j, t \mid y_j, t+\Delta t)\mathrm{d}y_j, \\
&\qquad = \frac{\partial P_i}{\partial y_j} \lim_{\Delta t \to 0} \frac{1}{\Delta t} \mathrm{E}[z_j - y_j] \\
&\qquad = 0,
\end{aligned}$$

since the expected amount of change under Brownian motion iz zero, *i.e.*, $\mathrm{E}[z_j - y_j] = 0$.

By a similar expansion, the second term of eq. (S.9) becomes:

$$\begin{aligned}
&\lim_{\Delta t \to 0} \frac{1}{\Delta t} \int_{z \in \mathbb{R}^c} f(\boldsymbol{z}, t \mid \boldsymbol{y}, t + \Delta t) \frac{1}{2} (\boldsymbol{z} - \boldsymbol{y}) \cdot \boldsymbol{H}_{P_i}(\boldsymbol{y}) \cdot (\boldsymbol{z} - \boldsymbol{y}) \mathrm{d}\boldsymbol{z} \\
&\qquad = \sum_{k=1}^{c} \sum_{l=1}^{c} \lim_{\Delta t \to 0} \frac{1}{2} \frac{1}{\Delta t} \int_{z \in \mathbb{R}^c} f(\boldsymbol{z}, t \mid \boldsymbol{y}, t + \Delta t) \frac{(z_k - y_k)(z_l - y_l)}{2} \frac{\partial^2 P_i}{\partial y_k y_l} \mathrm{d}\boldsymbol{z} \\
&\qquad = \sum_{k=1}^{c} \sum_{l=1}^{c} \frac{1}{2} \frac{\partial^2 P_i}{\partial y_k y_l} \lim_{\Delta t \to 0} \frac{1}{\Delta t} \int_{z \in \mathbb{R}^c} f(\boldsymbol{z}, t \mid \boldsymbol{y}, t + \Delta t) (z_k - y_k)(z_l - y_l) \mathrm{d}\boldsymbol{z}.
\end{aligned} \tag{S.12}$$

As before, we can rewrite the joint transition probability density of $z_l$ and $z_k$, marginal with the respect to the remaining characters, as

$$f(z_l, z_k, t \mid y_l, y_k, t + \Delta t) = \int_{z'} f(\boldsymbol{z}, t \mid \boldsymbol{y}, t + \Delta t) \mathrm{d}\boldsymbol{z}',$$

so that

$$\begin{aligned}
&\lim_{\Delta t \to 0} \frac{1}{\Delta t} \int_{z \in \mathbb{R}^c} f(\boldsymbol{z}, t \mid \boldsymbol{y}, t + \Delta t) (z_k - y_k)(z_l - y_l) \mathrm{d}\boldsymbol{z} \\
&\qquad = \lim_{\Delta t \to 0} \int_{z_k} \int_{z_l} (z_k - y_k)(z_l - y_l) f(z_l, z_k, t \mid y_l, y_k, t + \Delta t) \int_{z'} f(\boldsymbol{z}, t \mid \boldsymbol{y}, t + \Delta t) \mathrm{d}\boldsymbol{z}' \mathrm{d}z_k \mathrm{d}z_l \\
&\qquad = \lim_{\Delta t \to 0} \frac{1}{\Delta t} \int_{z_k} \int_{z_l} (z_k - y_k)(z_l - y_l) f(z_l, z_k, t \mid y_l, y_k, t + \Delta t) \mathrm{d}z_k \mathrm{d}z_l \\
&\qquad = \lim_{\Delta t \to 0} \frac{1}{\Delta t} \mathrm{E}[(z_k - y_k)(z_l - y_l)] \\
&\qquad = \lim_{\Delta t \to 0} \frac{1}{\Delta t} \mathrm{Cov}(z_k, z_l),
\end{aligned}$$

where the instantaneous covariance between $z_k$ and $z_l$, $\mathrm{Cov}(z_k, z_l)$, is defined as $\Sigma_{kl}$. Equation (S.12) becomes

$$\lim_{\Delta t \to 0} \frac{1}{\Delta t} \int_{z \in \mathbb{R}^c} f(\boldsymbol{z}, t \mid \boldsymbol{y}, t + \Delta t) \frac{1}{2} (\boldsymbol{z} - \boldsymbol{y}) \cdot \boldsymbol{H}_{P_i}(\boldsymbol{y}) \cdot (\boldsymbol{z} - \boldsymbol{y}) \mathrm{d}\boldsymbol{z} = \sum_{k=1}^{c} \sum_{l=1}^{c} \frac{\Sigma_{kl}}{2} \frac{\partial^2 P_i(\boldsymbol{y}, t)}{\partial y_k y_l}.$$

Following the definition of a Brownian motion process (Allen 2010), we assume that the probability of large jumps shrinks to zero as $\Delta t \to 0$; formally,

$$\lim_{\Delta t \to 0} \frac{1}{\Delta t} \int_{z \in \mathbb{R}^c} \prod_{m=1}^{c} f(\boldsymbol{z}, t \mid \boldsymbol{y}, t + \Delta t)(z_m - y_m)^{k_m} \mathrm{d}\boldsymbol{z} = 0, \ \sum_{m=1}^{c} k_m > 2.$$

This assumption allows the final term of eq. (S.9) to disappear. Ultimately, eq. (S.9) becomes

$$\lim_{\Delta t \to 0} \frac{1}{\Delta t} \int_{z \in \mathbb{R}^c} f(\boldsymbol{z}, t \mid \boldsymbol{y}, t + \Delta t)(P_i(\boldsymbol{z}, t) - P_i(\boldsymbol{y}, t)) \mathrm{d}\boldsymbol{z} = \sum_{k=1}^{c} \sum_{l=1}^{c} \frac{\Sigma_{kl}^i}{2} \frac{\partial^2 P_i(\boldsymbol{y}, t)}{\partial y_k y_l}, \tag{S.13}$$

where $\Sigma^i$ is the variance-covariance matrix of the $i^{\text{th}}$ discrete state.

##### S.1.4 Partial Differential Equation for the Transition Probability Density

Substitution equations (S.7), (S.8), and (S.13) into eq. (S.5) results in a set of partial differential equations, one per discrete state, $i$:

$$\frac{\partial P_i(\boldsymbol{y},t)}{\partial t} = \sum_{j \neq i}^{d} q_{ji} P_j(\boldsymbol{y},t) - \sum_{j \neq i}^{d} q_{ij} P_i(\boldsymbol{y},t) + \sum_{k=1}^{c} \sum_{l=1}^{c} \frac{\Sigma_{kl}^{i}}{2} \frac{\partial^2 P_i(\boldsymbol{y},t)}{\partial y_k y_l}. \tag{S.14}$$

These sets of partial differential equations have an intuitive interpretation: the first term represents the flow of probability *into* discrete state $i$, the second term represents the flow of probability *away from* discrete state $i$, and the third term represents the flow of probability between the continuous states within discrete state $i$.

#### S.2 Identifiability of the State-Dependent Model

To prove that a model is nonidentifiable, it is sufficient to find two combinations of parameters, $\theta$ and $\theta'$, that result in identical likelihoods. For the state-dependent model with background variation, we can demonstrate nonidentifiability if we can find $\theta$ and $\theta'$ that result in identical overall variance-covariance matrices, $\Sigma_l = \Sigma'_l$, for each lineage $l$.

We recall that the evolutionary variance-covariance matrix for lineage $l$ is a weighted sum of the state-dependent variance-covariance matrices:

$$\Sigma_l = \beta_l^2 \left[ \tau(\kappa_l, 0)\Sigma^0 + \tau(\kappa_l, 1)\Sigma^1 \right], \tag{S.15}$$

where

$$\Sigma^i = \zeta_i^2 \Sigma \tag{S.16}$$

is the state-dependent variance-covariance matrix in discrete state $i$ (see Main Text for details).

By rearranging equations (S.15) and (S.16), we see that $\Sigma_l$ is a scalar multiple of $\Sigma$:

$$\Sigma_l = \beta_l^2 \left[ \tau(\kappa_l, 0)\zeta_0^2 + \tau(\kappa_l, 1)\zeta_1^2 \right] \Sigma, \tag{S.17}$$

where the scalar, $\beta_l^2 \left[ \tau(\kappa_l, 0)\zeta_0^2 + \tau(\kappa_l, 1)\zeta_1^2 \right]$, is a function of the various sources of rate variation. By substituting (S.17) into the conditions for nonidentifiability, $\Sigma_l = \Sigma'_l$, we get

$$\beta_l^2 \left[ \tau(\kappa_l, 0)\zeta_0^2 + \tau(\kappa_l, 1)\zeta_1^2 \right] = \beta_l'^2 \left[ \tau(\kappa_l, 0)\zeta_0'^2 + \tau(\kappa_l, 1)\zeta_1'^2 \right].$$

If we allow $\zeta_0'^2 = \zeta_1'^2 = 1$, then

$$\beta_l^2 \left[ \tau(\kappa_l, 0)\zeta_0^2 + \tau(\kappa_l, 1)\zeta_1^2 \right] = \beta_l'^2 t_l,$$

since $\tau(\kappa_l, 0) + \tau(\kappa_l, 1) = t_l$ (the branch length). Because $\beta_l^2$ and $\beta_l'^2$ are unconstrained, it is clear that we can find some values of $\theta$ and $\theta'$ that have identical likelihoods. Therefore, the state-dependent model with background-rate variation is nonidentifiable.

#### S.3 Numerical Consequences of Sequential Analysis

Our implementation of the discrete-state-dependent continuous-character model uses data augmentation to simultaneously sample discrete-character histories and the parameters of the state-dependent model during MCMC; certainly, this is not the only possible solution. However, we show that another MCMC solution—a "sequential analysis"—may fail in certain circumstances, and that those circumstances are actually related to the biological question that motivates the model.

For the data-augmented model, the posterior distribution of the continuous-character model parameters, $\theta_c$, the discrete-character histories, $\boldsymbol{\kappa}$, and the discrete-character model parameters, $\theta_d$, given the data, $\mathcal{X}$ and $\mathcal{Y}$, and the phylogeny, $\Psi$, is:

$$P(\theta_c, \theta_d, \boldsymbol{\kappa} \mid \mathcal{X}, \mathcal{Y}, \Psi) \propto P(\mathcal{X}, \mathcal{Y}, \boldsymbol{\kappa} \mid \theta_c, \theta_d, \Psi) P(\theta_c) P(\theta_d) \tag{S.18}$$

$$\propto P(\mathcal{X}, \boldsymbol{\kappa} \mid \theta_d, \Psi) P(\mathcal{Y} \mid \boldsymbol{\kappa}, \theta_c, \Psi) P(\theta_c) P(\theta_d). \tag{S.19}$$

A theoretically equivalent solution involves performing a "sequential" Bayesian analysis (Caetano and Harmon 2017, 2018). Under this sequential procedure, we would first generate a posterior distribution of discrete-character histories:

$$P(\boldsymbol{\kappa}, \theta_d \mid \mathcal{X}, \Psi) \propto P(\mathcal{X}, \boldsymbol{\kappa} \mid \theta_d, \Psi) P(\theta_d), \tag{S.20}$$

and then use the marginal posterior distribution of $\boldsymbol{\kappa}$ as a prior distribution in a second analysis:

$$P(\theta_c, \boldsymbol{\kappa} \mid \mathcal{Y}, \Psi) \propto P(\mathcal{Y} \mid \boldsymbol{\kappa}, \theta_c, \Psi) P(\boldsymbol{\kappa}) P(\theta_c). \tag{S.21}$$

In other words, this approach involves using the samples from the first step as a "numerical" prior distribution for the second step. In the second step, the MCMC is initialized with a random draw from the input (numerical-prior) distribution; the MCMC then proposes changes to the discrete-character history by drawing a sample (at random) from the input distribution, and accepting the proposal with probability:

$$A = \min \left[ 1, \frac{P(\mathcal{Y} \mid \boldsymbol{\kappa}', \theta_c, \Psi)}{P(\mathcal{Y} \mid \boldsymbol{\kappa}, \theta_c, \Psi)} \times \frac{P(\boldsymbol{\kappa}')}{P(\boldsymbol{\kappa})} \times \frac{f(\boldsymbol{\kappa})}{f(\boldsymbol{\kappa}')} \right]. \tag{S.22}$$

Because new states, $\boldsymbol{\kappa}'$, are proposed by drawing randomly from the numerical sample, the last two terms of the acceptance probability cancel out. Combined with proposals on $\theta_c$, this MCMC should—in principle—converge to the joint posterior distribution, eq. (S.21).

We can see that eq. (S.21) is equivalent to eq. (S.19) by noting that each sample of $\boldsymbol{\kappa}$ in the posterior of the first step is associated with a sample of $\theta_d$, *i.e.*, the first step samples the joint posterior distribution of character histories and discrete-character model parameters. We can therefore "reconstruct" the joint posterior of $\theta_c$, $\theta_d$, and $\boldsymbol{\kappa}$ from the sequential analysis by "looking up" the value of $\theta_d$ associated with each sample of $\boldsymbol{\kappa}$ in the posterior distribution generated by the second step. In other words, we are free to rewrite eq. (S.21) as:

$$P(\theta_c, \theta_d, \boldsymbol{\kappa} \mid \mathcal{Y}, \Psi) \propto P(\mathcal{Y} \mid \boldsymbol{\kappa}, \theta_c, \Psi) P(\boldsymbol{\kappa}, \theta_d) P(\theta_c). \tag{S.23}$$

Substituting eq. (S.20) into eq. (S.23) results in:

$$P(\theta_c, \theta_d, \boldsymbol{\kappa} \mid \mathcal{X}, \mathcal{Y}, \Psi) \propto P(\mathcal{Y} \mid \boldsymbol{\kappa}, \theta_c, \Psi) P(\mathcal{X}, \boldsymbol{\kappa} \mid \theta_d, \Psi) P(\theta_d) P(\theta_c), \tag{S.24}$$

which is obviously equivalent to eq. (S.19).

Although the joint (data-augmentation) and sequential analyses are theoretically equivalent, they may behave quite differently in practice. The sequential analysis requires that we first generate a posterior distribution of $\boldsymbol{\kappa}$ based solely on the discrete-character data, $\mathcal{X}$. As this sample is a numerical approximation of a probability distribution, it effectively assigns zero probability density to all unsampled character histories. If the joint posterior distribution, $P(\theta_c, \theta_d, \boldsymbol{\kappa} \mid \mathcal{X}, \mathcal{Y}, \Psi)$, assigns high posterior density to character histories that have low (*i.e.*, 0) posterior density in the first step of the sequential analysis, then the second step of the sequential analysis will be unable to reconstruct the joint posterior distribution.

The degree to which the full joint posterior distribution and the sample of character histories differ depends on the amount of information the continuous characters contain about the discrete-character history; the more $\mathcal{Y}$ depends on $\boldsymbol{\kappa}$, the worse this problem should be. That is, the problem grows worse as the rates of continuous-character evolution depend more strongly on the state of the discrete character, which is exactly the scenario of interest! The effect will also clearly depend on the number of samples generated in the first step: with an infinite sample, no character histories would have zero prior density, so the problem can—in principle—be overcome by generating a sufficiently large numerical sample. The number of samples required to provide a sufficient approximation will naturally depend on the dataset, but intuitively we can see that we will need more samples as the disparity between the distributions increases.

We conducted experiments with the haemulid dataset to demonstrate the potential pitfalls of sequential analysis for the state-dependent model. We motivate this by first showing the posterior distribution of $k$ (the number of transitions in the character history) under a discrete-character model compared to the posterior distribution of $k$ under the state-dependent multivariate Brownian motion model without background-rate variation (Fig. S1). In this example, it is clear that the posterior distributions of character histories are extremely different, which should have drastic consequences for the sequential analysis.

We performed a sequential analysis by first generating the posterior distribution of character histories under the discrete-character model using eq. (S.20). We drew 1000 samples from the posterior distribution to form the numerical prior for the second step of the sequential analysis. In the second step, we estimated the posterior distribution of the state-dependent model without background-rate variation using eq. (S.21) and the proposal mechanism described above. We performed four inde-

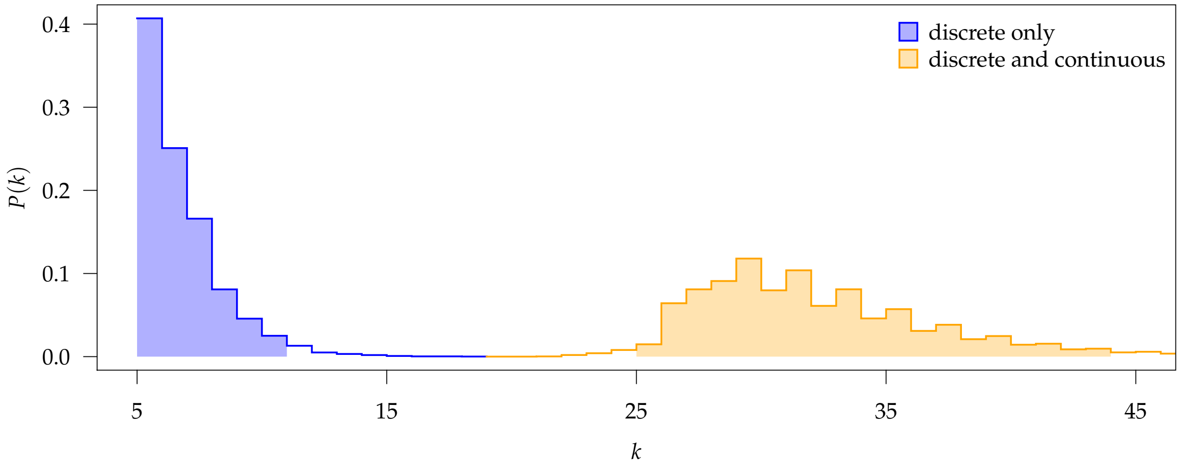


**Figure S1:** The marginal posterior distribution of the number of transitions, $k$, when only analyzing the discrete-characters (blue distribution, with 95% CI shaded) and when analyzing the discrete- and continuous-characters jointly under the state-dependent model (orange distribution, with 95% CI shaded).

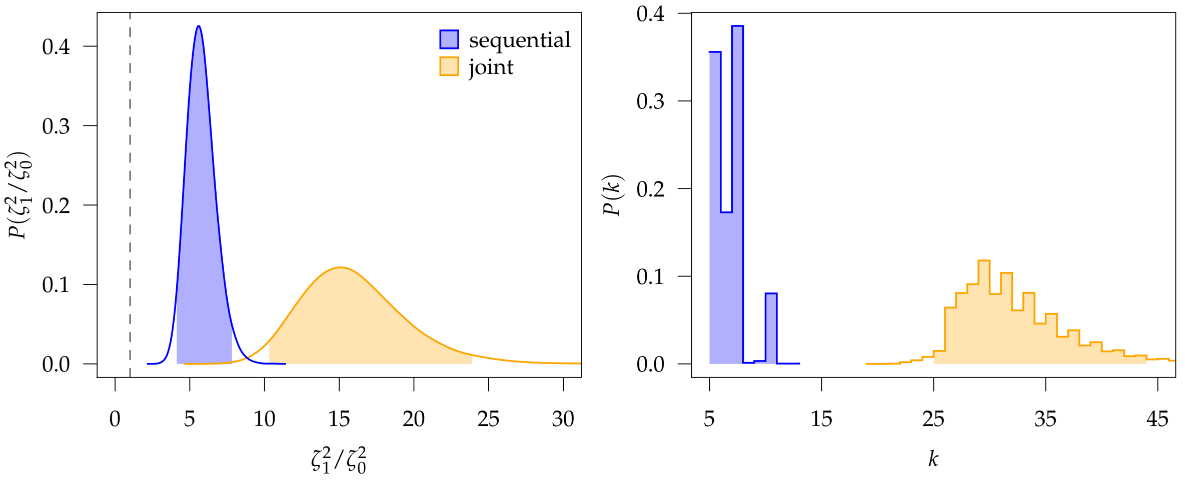


**Figure S2:** The marginal posterior distributions from a "sequential" MCMC analysis (blue distribution, with 95% CI shaded) and a simultaneous analysis using data augmentation (orange distribution, with 95% CI shaded). The left panel depicts distributions of the focal parameters, $\zeta_1^2/\zeta_0^2$; the dashed vertical line corresponds to $\zeta_1^2/\zeta_0^2 = 1$. The right panel depicts distributions of the number of transition events, $k$.

pendent Metropolis-coupled MCMC replicates for the sequential analysis, and confirmed that they converged to the same distribution. The resulting marginal posterior density of the focal parameter, $\zeta_1^2/\zeta_0^2$, inferred under the sequential procedure differs substantially from that obtained from analyses under the joint (data augmentation) approach (Fig. S2, left panel). Moreover, the number of transition events inferred under these two approaches is similarly distorted (Fig. S2, right panel). This behavior implies that—although the sequential MCMC analyses converged to *a distribution*—they did not converge to *the joint posterior distribution*, and the resulting samples cannot be interpreted as being drawn from the joint posterior probability distribution.

We note that the state-dependent model without background-rate variation provides very unrealistic estimates of the state-dependent rates of continuous-character evolution and of the history of discrete-character evolution (see Main Text). In this light, it may seem that the sequential analysis has "protected" us from drawing unrealistic inferences about state-dependent rates of continuous-character evolution, or that the sequential analysis is "conservative" with respect to our hypothesis. However, whether a model provides an adequate description of reality and whether we can reliably estimate parameters under that model are distinct issues, and it is probably unsafe to rely on numerical pathologies to solve modeling inadequacies.

A similar issue may apply to related methods that use Monte Carlo simulation to integrate over discrete-character histories (*e.g.*, Mayrose and Otto 2010). We begin by describing these approaches and demonstrating that they are equivalent to data augmentation when implemented in a Bayesian setting. Integrating the posterior distribution of the data-augmented model, eq. (S.19), over the space of character histories ($\boldsymbol{K}$) yields the posterior distribution of the continuous- and discrete-model parameters:

$$P(\theta_c, \theta_d \mid \mathcal{X}, \mathcal{Y}, \Psi) = \int_{\boldsymbol{K}} P(\theta_c \mid \boldsymbol{\kappa}, \mathcal{Y}, \Psi) P(\boldsymbol{\kappa} \mid \theta_d, \mathcal{X}, \Psi) \, \mathrm{d}\boldsymbol{\kappa} \tag{S.25}$$

$$\propto \left[ \int_{\boldsymbol{K}} P(\mathcal{X}, \boldsymbol{\kappa} \mid \theta_d, \Psi) P(\mathcal{Y} \mid \boldsymbol{\kappa}, \theta_c, \Psi) \, \mathrm{d}\boldsymbol{\kappa} \right] P(\theta_c, \theta_d), \tag{S.26}$$

where eq. (S.26) results from substituting the right-hand side of eq. (S.19) into eq. (S.25). Expanding the joint probability of the discrete data and character histories,

$$P(\theta_c, \theta_d \mid \mathcal{X}, \mathcal{Y}, \Psi) \propto \left[ \int_{\boldsymbol{K}} P(\mathcal{X} \mid \theta_d, \Psi) P(\boldsymbol{\kappa} \mid \mathcal{X}, \theta_d, \Psi) P(\mathcal{Y} \mid \boldsymbol{\kappa}, \theta_c, \Psi) \, \mathrm{d}\boldsymbol{\kappa} \right] P(\theta_c, \theta_d) \tag{S.27}$$

$$\propto \underbrace{\left[ P(\mathcal{X} \mid \theta_d, \Psi) \int_{\boldsymbol{K}} P(\boldsymbol{\kappa} \mid \mathcal{X}, \theta_d, \Psi) P(\mathcal{Y} \mid \boldsymbol{\kappa}, \theta_c, \Psi) \, \mathrm{d}\boldsymbol{\kappa} \right]}_{\text{joint probability of } \mathcal{X}, \mathcal{Y}} P(\theta_c, \theta_d). \tag{S.28}$$

Monte Carlo integration computes the joint probability of $\mathcal{X}$ and $\mathcal{Y}$ by averaging over $n$ samples of $\boldsymbol{\kappa}$ drawn from its conditional distribution, $P(\boldsymbol{\kappa} \mid \mathcal{X}, \theta_d, \Psi)$:

$$P(\mathcal{X} \mid \theta_d, \Psi) \int_{\boldsymbol{K}} P(\boldsymbol{\kappa} \mid \mathcal{X}, \theta_d, \Psi) P(\mathcal{Y} \mid \boldsymbol{\kappa}, \theta_c, \Psi) \, \mathrm{d}\boldsymbol{\kappa} \approx \frac{P(\mathcal{X} \mid \theta_d, \Psi)}{n} \sum_{i=1}^{n} P(\mathcal{Y} \mid \boldsymbol{\kappa}_i, \theta_c, \Psi) \tag{S.29}$$

We identify the joint probability of $\mathcal{X}$ and $\mathcal{Y}$ in eq. (S.28) to make it clear that this term does not depend on our Bayesian formulation (as it does not contain any prior distributions): this term also describes how to compute the likelihood under a maximum-likelihood approach (*c.f.*, equation 4 in Mayrose and Otto 2010). In a Bayesian context, Monte Carlo integration theoretically provides the same posterior distribution as data augmentation after the discrete-character histories are marginalized out (*c.f.*, eq. [S.25]). However, Monte Carlo integration—like the sequential MCMC approach—may fail if the distribution used to generate the discrete-character histories, $\boldsymbol{\kappa_i} \sim P(\boldsymbol{\kappa} \mid \mathcal{X}, \theta_d, \Psi)$, assigns approximately zero probability to character histories that have high joint probability, $P(\mathcal{X}, \mathcal{Y}, \boldsymbol{\kappa} \mid \theta_c, \theta_d, \Psi)$.

We emphasize that the sequential approach and Monte Carlo integration are both theoretically valid solutions to the problem of computing the joint probability of $\mathcal{X}$ and $\mathcal{Y}$, and we have no reason to believe that they are *generally* problematic. However, we have demonstrated that may *sometimes* be problematic, and therefore believe the practical behavior of these approaches deserves further study. Accordingly, empirical applications of these approaches would ideally explore the impact of the number of samples used to approximate the prior distribution—and/or the number of Monte Carlo replicates used to integrate the joint probability—on estimates of the model parameters. However, even this "best practice" could fail to detect pathologies in cases (such as the example considered above) where there is considerable disparity between the distribution used to sample character histories and the target distribution.

#### S.4 Standard Deviation of the Lognormal Distribution

We introduce a constant, $H$, which is the standard deviation of a lognormal random variable, $X$, such that the 95% probability interval of the random variable ranges over one order of magnitude.

We seek to find a value of $\sigma$ such that:

$$Q(0.975) = 10Q(0.025), \tag{S.30}$$

where $Q(p)$ is the quantile function of a lognormal distribution evaluated at $p$:

$$Q(p) = \mathrm{e}^{\mu+\sigma\Phi^{-1}(p)}, \tag{S.31}$$

where $\Phi^{-1}$ is the quantile function of the standard normal distribution.

Combining (S.30) and (S.31) and solving for $\sigma$:

$$\sigma = \frac{\ln 10}{\Phi^{-1}(0.975) - \Phi^{-1}(0.025)} \tag{S.32}$$

$$\sigma \approx 0.587405 \tag{S.33}$$

$$H = 0.587405. \tag{S.34}$$

#### S.5 MCMC Proposals

##### S.5.1 Reversible-jump proposals for symmetric and asymmetric models

We have specified a mixture model as a prior on the discrete-character transition rates, $\boldsymbol{q}$. We use the transdimensional generalization of the Metropolis-Hastings algorithm, reversible-jump MCMC (Green 1995), to propose changes between symmetric and asymmetric models. If the chain is visiting a symmetric model, we propose to move to an asymmetric model by drawing new values of $\boldsymbol{q}$ from the prior distribution, Dirichlet($\alpha_q$). The acceptance probability for this proposal is:

$$A = \min\left[1, \frac{P(\mathcal{X},\mathcal{Y},\boldsymbol{\kappa} \mid \theta')}{P(\mathcal{X},\mathcal{Y},\boldsymbol{\kappa} \mid \theta)} \times \frac{P(\text{asym})}{P(\text{sym})} \times \frac{P_a(\boldsymbol{q}')}{P_s(\boldsymbol{q})} \times \frac{f(\boldsymbol{q})}{f(\boldsymbol{q}')} \times |J|\right], \tag{S.35}$$

where $P(\text{sym})$ and $P(\text{asym})$ are the mixture probabilities of the symmetric and asymmetric models, $P_s$ and $P_a$ refer to the prior distribution of the symmetric and asymmetric models, and $J$ is the determinant of the Jacobian of transformation between the proposed and current states of the Markov chain. Because we draw the new value, $\boldsymbol{q}'$, from the asymmetric prior distribution,

$$\frac{f(\boldsymbol{q})}{f(\boldsymbol{q}')} = \frac{1}{P_a(\boldsymbol{q})} \tag{S.36}$$

$$|J| = 1, \tag{S.37}$$

the acceptance probability becomes:

$$A = \min\left[1, \frac{P(\mathcal{X},\mathcal{Y},\boldsymbol{\kappa} \mid \theta')}{P(\mathcal{X},\mathcal{Y},\boldsymbol{\kappa} \mid \theta)} \times \frac{P(\text{asym})}{P(\text{sym})} \times \frac{P_a(\boldsymbol{q}')}{P_s(\boldsymbol{q})} \times \frac{1}{P_a(\boldsymbol{q}')}\right]. \tag{S.38}$$

To propose a change from an asymmetric model to a symmetric model, we set $q_{01} = q_{10}$, and accept with probability:

$$A = \min\left[1, \frac{P(\mathcal{X},\mathcal{Y},\boldsymbol{\kappa} \mid \theta')}{P(\mathcal{X},\mathcal{Y},\boldsymbol{\kappa} \mid \theta)} \times \frac{P(\text{sym})}{P(\text{asym})} \times \frac{P_s(\boldsymbol{q}')}{P_a(\boldsymbol{q})} \times \frac{P_a(\boldsymbol{q})}{1}\right]. \tag{S.39}$$

##### S.5.2 Proposals on discrete-character histories

To propose a change to the discrete-character histories, we draw a new value of $\boldsymbol{\kappa}$, and accept the proposed state with probability:

$$A = \min\left[1, \frac{P(\mathcal{X},\mathcal{Y},\boldsymbol{\kappa}' \mid \theta)}{P(\mathcal{X},\mathcal{Y},\boldsymbol{\kappa} \mid \theta)} \times \frac{\cancel{P(\theta)}}{\cancel{P(\theta)}} \times \frac{f(\boldsymbol{\kappa})}{f(\boldsymbol{\kappa}')}\right], \tag{S.40}$$

where the prior ratio cancels out because the parameters are unchanged. Expanding the augmented likelihood, we get

$$A = \min\left[1, \frac{P(\mathcal{Y} \mid \boldsymbol{\kappa}',\theta)}{P(\mathcal{Y} \mid \boldsymbol{\kappa},\theta)} \times \frac{P(\mathcal{X},\boldsymbol{\kappa}' \mid \theta)}{P(\mathcal{X},\boldsymbol{\kappa} \mid \theta)} \times \frac{f(\boldsymbol{\kappa})}{f(\boldsymbol{\kappa}')}\right]. \tag{S.41}$$

We propose new values of $\boldsymbol{\kappa}$ by simulating new character histories in proportion to their probability given the data and the discrete-character model, $Q$. Specifically, we choose a node in the tree at random, then sample a new discrete state at that node in proportion to its probability, given the states at the incident nodes and the rate matrix, $Q$. We then simulate character histories along the incident branches, and reject those histories that end in discrete states that do not match the discrete states of the internal nodes. This procedure is based on the stochastic-mapping algorithm (Nielsen 2002; Huelsenbeck et al. 2003), and ensures that proposed values of the character histories have high posterior probabilities. The acceptance ratio for this move is:

$$A = \min\left[1, \frac{P(\mathcal{Y} \mid \boldsymbol{\kappa}', \theta)}{P(\mathcal{Y} \mid \boldsymbol{\kappa}, \theta)} \times \frac{\cancel{P(\mathcal{X}, \boldsymbol{\kappa}' \mid \theta)}}{\cancel{P(\mathcal{X}, \boldsymbol{\kappa} \mid \theta)}} \times \frac{\cancel{P(\mathcal{X}, \boldsymbol{\kappa} \mid \theta)}}{\cancel{P(\mathcal{X}, \boldsymbol{\kappa}' \mid \theta)}}\right]. \tag{S.42}$$

The proposal ratio and the discrete-character component of the augmented-likelihood ratio cancel out, so that the new character histories are accepted with probability proportional to the ratio of the conditional densities of the continuous characters.

#### S.6 Empirical Analyses

##### S.6.1 MCMC Diagnosis

We performed four independent MCMC analyses for each of the models described in the sections below. We performed all analyses in `RevBayes` (Höhna et al. 2016). We provide details on the specific proposals, proposal weights, and MCMC settings within the scripts included in our supplementary-data archive.

We diagnosed each analysis by computing the effective sample size and Geweke's diagnostic (Geweke et al. 1991) for each parameter using the `R` package `coda` (Plummer et al. 2006); we considered an analysis a failure if more than 5% of the diagnostics failed (ESS $< 500$, Geweke's $p$-value $< 0.05$). We combined samples from the four independent (and successful) MCMC analyses for down-stream summaries.

##### S.6.2 Model Adequacy

We used posterior-predictive simulation to assess the adequacy of the substitution models used for the phylogenetic analyses described below. To do so, we simulated datasets from the joint posterior density of model parameters sampled by MCMC, and for each simulated dataset computed the log-multinomial test statistic (Bollback 2002), $T_x$, generating a posterior-predictive distribution of the statistic, conditional on the data, $\mathcal{S}$, $P(T_x \mid \mathcal{S})$. We then computed the observed test statistic (computed on the observed dataset), $T_x^{\text{obs}}$, and compared it to the predictive distribution; we consider a model to be inadequate if the observed statistic is not contained in the 95% credible interval of the posterior-predictive distribution (*i.e.*, the predictive interval).

##### S.6.3 Phylogenetic Analyses

We assembled a molecular dataset by subsampling the alignments from Tavera et al. (2018) to include only the 49 species that were represented in our morphological dataset. The resulting sequence dataset included three mitochondrial loci (*16S*, *COI*, and *CYTB*) and four nuclear loci (*RAG1*, *RAG2*, *S7*, and *TMO 4c4*). We first estimated the phylogeny and model parameters for each gene region separately in order to assess model adequacy. We then combined all genes into a (partitioned) concatenated analysis, and again assessed adequacy of the composite model. Finally, we estimated relative divergence times using a relaxed-clock model.

###### S.6.3.1 Individual-gene analyses

For each gene region, we estimated the posterior distribution of unrooted phylogenies under a GTR $+ \Gamma$ model of molecular evolution (partitioned by codon position for protein-coding genes, and by stem and loop regions for *16S*). We used uninformative priors on the substitution-model parameters (Zwickl and Holder 2004), a discrete-uniform prior on the tree topology, a flat Dirichlet prior on the relative-branch lengths, and a log-uniform prior on the tree length (the sum of the branch lengths). We provide more explicit details of our prior settings in the analysis scripts in the supplementary-data archive.

We assessed the adequacy for each data partition under the GTR $+ \Gamma$ model using posterior-predictive simulation as described above; none of the data partitions failed the posterior-predictive test (Fig. S3).

###### S.6.3.2 Concatenated analysis

We estimated the posterior distribution of unrooted phylogenies for the concatenated alignments that we partitioned by gene region, as well as by codon position or stem/loop regions (as appropriate for each gene region). We specified a composite substitution model comprising an independent GTR $+ \Gamma$ substitution model for each data subset, and specified priors as described for the individual-gene analyses (see supplementary-data archive for scripts).

We assessed the adequacy of the composite substitution model following the same procedures used for the individual-gene analyses; again, none of the data partitions failed the posterior-predictive test (Fig. S4).

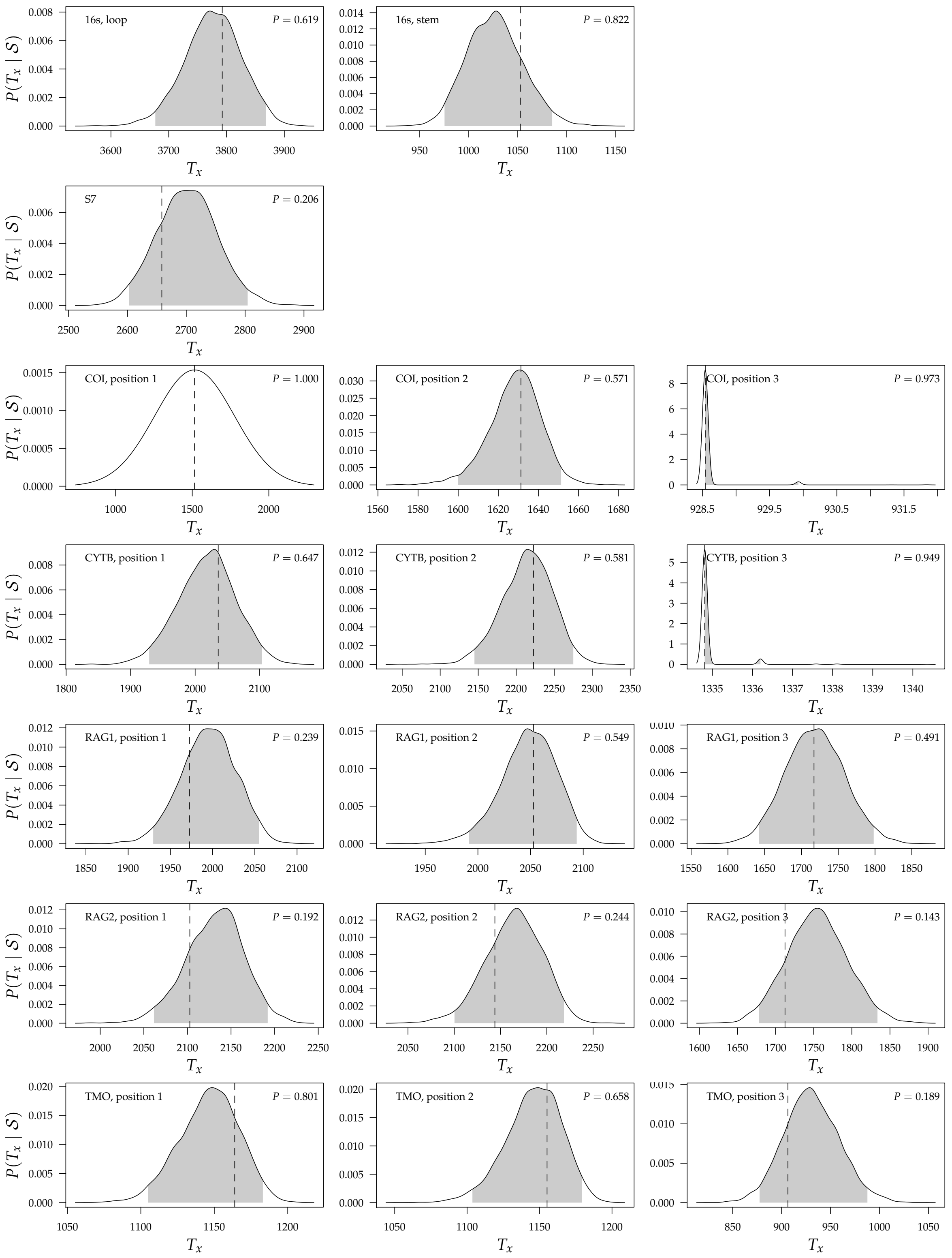


**Figure S3:** Posterior-predictive distributions for the gene-specific analyses. Each panel depicts the posterior-predictive distribution of the (log)-multinomial test statistic, $T_x$, for each data subset of a specific gene region (one gene region per row). Shaded regions indicate the 95% posterior-predictive interval; dashed vertical lines indicate the value of the test statistic for the observed data subset. Dashed lines that are contained by the shaded regions indicate that the model is adequate according to the multinomial likelihood test statistic. The posterior-predictive $p$-value (upper-right corner of each panel) is the fraction of simulated statistics that are smaller than or equal to the observed statistic.

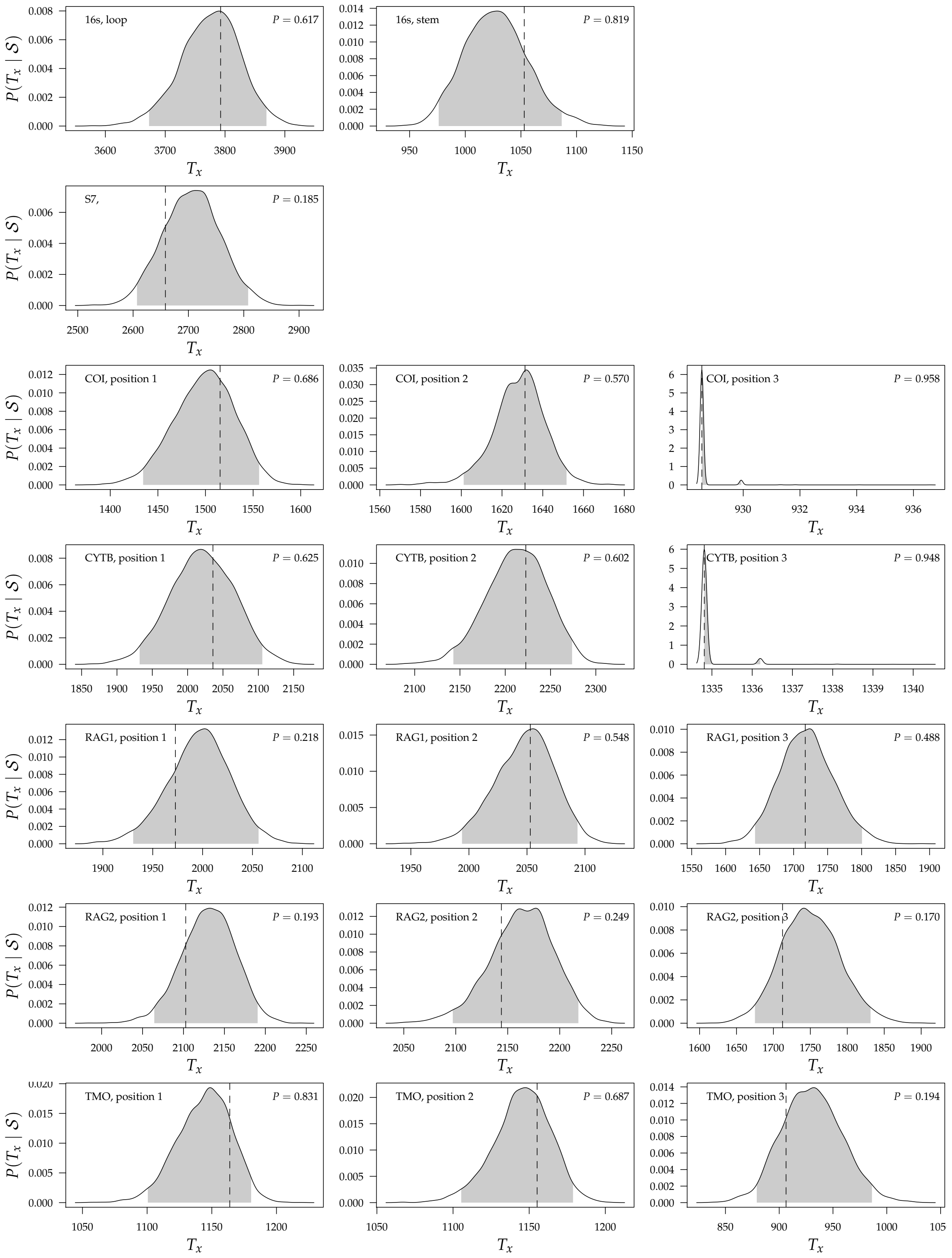


**Figure S4:** Posterior-predictive distributions for the concatenated analyses. Each panel depicts the posterior-predictive distribution of the (log)-multinomial test statistic, $T_x$, for each data subset of a specific gene region (one gene region per row). Shaded regions indicate the 95% posterior-predictive interval; dashed vertical lines indicate the value of the test statistic for the observed data subset. Dashed lines that are contained by the shaded regions indicate that the model is adequate according to the multinomial likelihood test statistic. The posterior-predictive *p*-value (upper-right corner of each panel) is the fraction of simulated statistics that are smaller than or equal to the observed statistic.

##### S.6.3.3 Divergence-time estimation

We estimated the posterior distribution of ultrametric phylogenies under a relaxed-clock model using the concatenated dataset. As with the previous analyses, we partitioned each gene region by codon position or stem/loop region, as appropriate. We used a constant-rate sampled birth–death process prior on the node ages, and an uncorrelated lognormal branch-rate prior model. Because we lack fossil calibrations for this group, and because we are only interested in *relative* rates of evolution, we constrained the age of the root to 1 arbitrary time unit. As before, we provide more explicit details of our prior settings in the analysis scripts included the supplementary-data archive. We computed the maximum *a posteriori* (MAP) chronogram (Figure S5) from the posterior distribution of ultrametric trees.

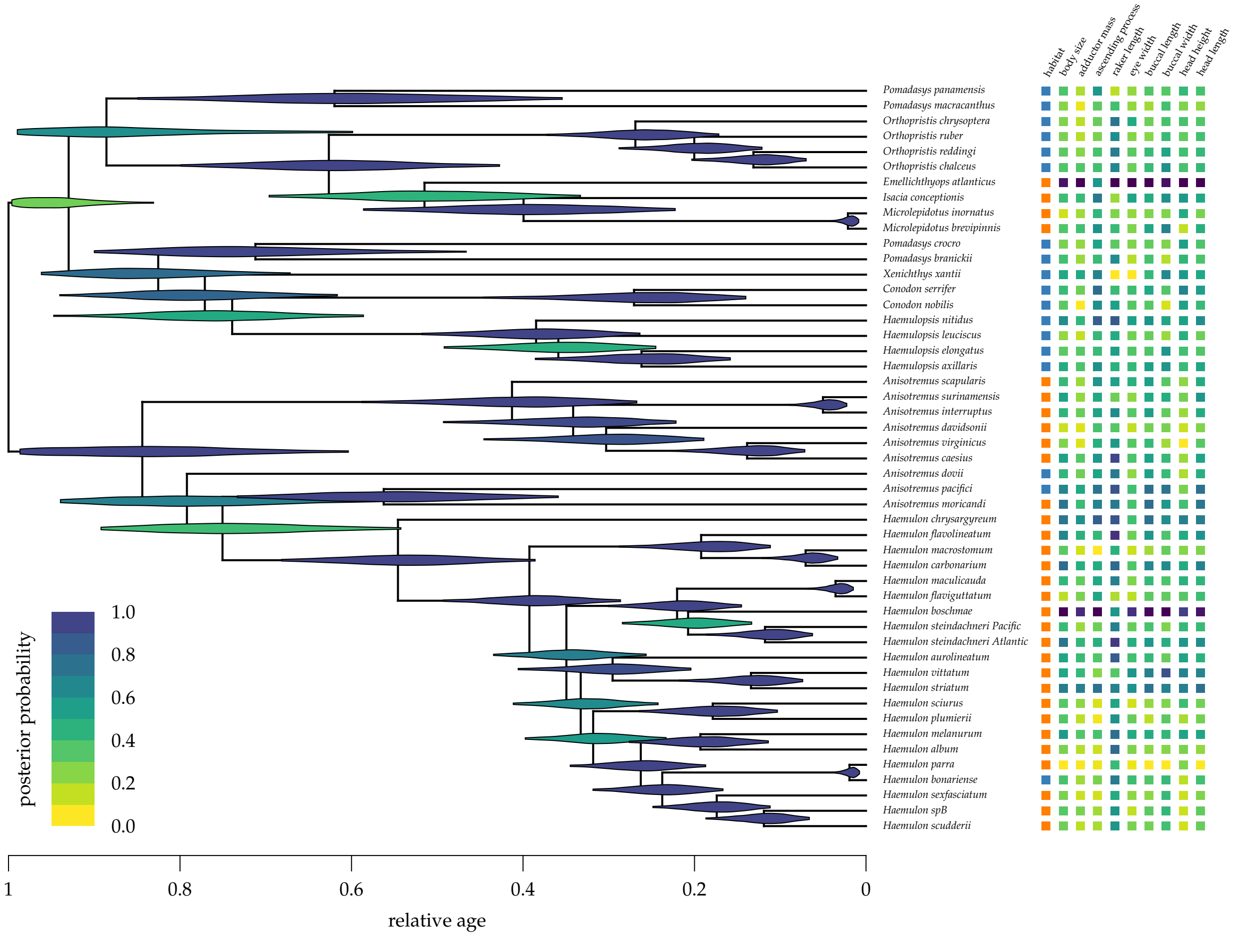


**Figure S5:** The estimated maximum *a posteriori* chronogram for haemulids. Violins represent the posterior densities for the relative age of each node truncated at their 95% CIs and colored by their posterior probability (see inset legend). At right, we indicate the trophic-character data for each species that we used to estimate the parameters of the state-dependent model: the discrete habitat is indicated in the first column (orange for reef-dwelling, blue for non-reef-dwelling) and the continuous characters are listed in the remaining columns (on a continuum from blue to yellow for small to large character values, respectively).

##### S.6.4 Joint Analysis of Phylogeny and Character Evolution

The empirical analyses of the state-dependent model that we describe in the Main Text (and in the sections below) assume that the MAP chronogram is a perfect estimate of the phylogeny and divergence times; because these analyses condition on the MAP tree, we refer to them as "conditional" analyses. However, it is also possible to estimate the parameters of the state-dependent model and the phylogeny simultaneously, which we refer to as "joint" analysis. The joint analysis naturally averages our estimates of the state-dependent model parameters over the posterior distribution of trees and divergence times, which, in principle, results in estimates that are robust to phylogenetic uncertainty.

We investigated the impact of phylogenetic uncertainty by simultaneously inferring the phylogeny (with divergence times) and the state-dependent model parameters from a combined dataset that included both the sequence alignments, $\mathcal{S}$ (describe above), as well as the comparative data, $\mathcal{X}$ and $\mathcal{Y}$. Our joint analyses estimated the joint posterior distribution of the phylogeny with divergence times, $\Psi$, the substitution-model parameters, $\theta$, and the parameters of the state-dependent model:

$$\begin{aligned} P(\boldsymbol{\zeta^2}, \boldsymbol{\beta^2}, \mu, \nu, \boldsymbol{\sigma^2}, R, Q, \boldsymbol{\kappa}, \theta, \Psi \mid \mathcal{X}, \mathcal{Y}, \mathcal{S}) \propto\ & P(\mathcal{X}, \boldsymbol{\kappa} \mid Q, \Psi) P(\mathcal{Y} \mid \boldsymbol{\zeta^2}, \boldsymbol{\beta^2}, \boldsymbol{\sigma^2}, R, \boldsymbol{\kappa}, \Psi) \\ & P(\mathcal{S} \mid \Psi, \theta) P(\Psi \mid \theta) \\ & P(\boldsymbol{\zeta^2}, \boldsymbol{\beta^2}, \mu, \nu, \boldsymbol{\sigma^2}, R, Q, \theta). \end{aligned}$$

We used the same data subsets, composite substitution model, and priors as those used in our divergence-time analyses, above (see supplementary-data archive).

We compared the posterior distribution of the state-dependent rate-ratio, $\zeta_1^2 / \zeta_0^2$, between the conditional analysis and the joint analysis (Fig. S6). It appears that, in this case, the posterior estimate of the focal parameter is not strongly influenced by phylogenetic uncertainty.

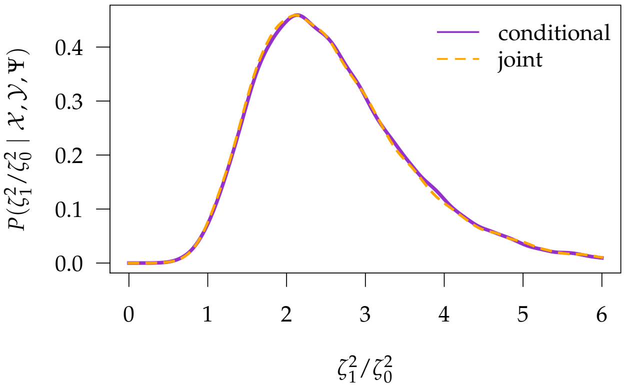


**Figure S6:** The marginal posterior density of the state-dependent rate-ratio, $\zeta_1^2\ \zeta_0^2$, inferred from the MAP tree (*conditional*, blue density) and incorporating phylogenetic uncertainty (*joint*, orange density).

##### S.6.5 Prior Sensitivity

We assessed the sensitivity of posterior estimates to the choice of priors by performing analyses of the haemulid data on the fixed (MAP) tree under various prior settings. For each parameter, we varied the prior over a set of values, while fixing the priors for the remaining parameters to their "default" values, as described in the *Priors* section of the Main Text. In this section, we describe the set of priors that we explored for each parameter, and discuss their effects on estimates of the corresponding parameter and also on estimates of the "focal parameter", *i.e.*, the state-dependent rate ratio, $\zeta_1^2/\zeta_0^2$.

###### S.6.5.1 State-dependent rates

We varied the prior on the state-dependent relative rates, $\alpha_{\zeta^2}$, over five values: $\alpha_{\zeta^2} \in \{1/4, 1/2, 1, 2, 4\}$. These prior densities, and the corresponding marginal posterior densities, are depicted in Fig. S7. The marginal posterior densities of the relative-rate ratio, $\zeta_1^2/\zeta_0^1$, are depicted in Fig. S8. (Because $\boldsymbol{\zeta}^2/2$ is a vector of length 2, we plot the prior and posterior distributions of the first element, $\zeta_0^2/2$.)

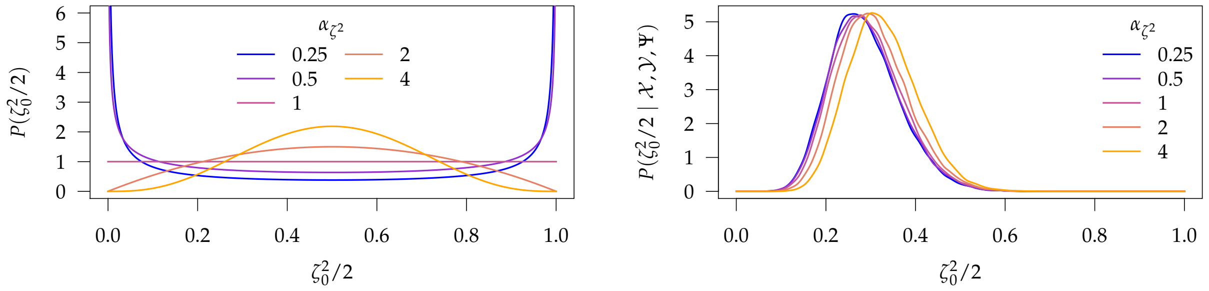


**Figure S7:** Prior sensitivity experiment for the state-dependent relative-rate parameter, $\zeta_0^2/2$. Left panel: the five prior probability densities. Right panel: the five corresponding marginal posterior probability densities.

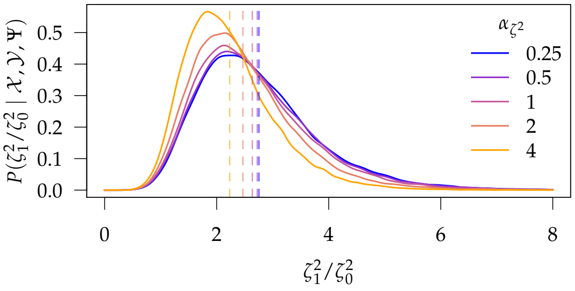


**Figure S8:** Marginal posterior probability densities of the relative-rate ratio, $\zeta_1^2/\zeta_0^1$, for each of the five priors on the state-dependent relative-rate parameter, $\zeta^2/2$. The dashed horizontal lines are the posterior means for each of the five marginal posterior densities.

##### S.6.5.2 Character-specific rates

We varied the prior on the character-specific relative-rate parameters, $\alpha_{\sigma^2}$, over five values: $\alpha_{\sigma^2} \in \{1/4, 1/2, 1, 2, 4\}$. These prior densities, and the corresponding marginal posterior densities, are depicted in Figs. S9 and S10, respectively. The marginal posterior densities of the relative-rate ratio, $\zeta_1^2/\zeta_0^1$, are depicted in Fig. S11. (Because $\boldsymbol{\sigma}^2/c$ is a vector of length $c$, we plot the prior distribution of the $i^{\text{th}}$ element, $\sigma_i^2/c$.)

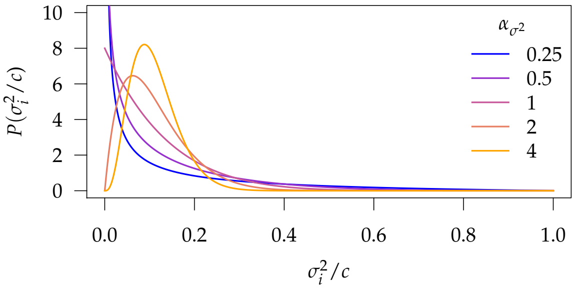


**Figure S9:** Marginal prior probability probability densities for the character-specific relative-rate parameters, $\sigma_i^2/c$.

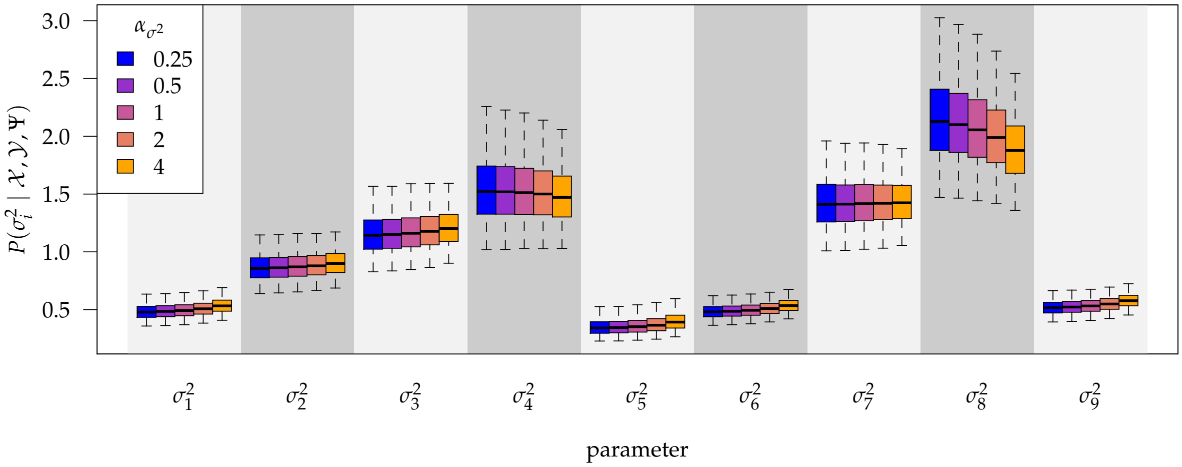


**Figure S10:** Boxplots of the marginal posterior probability densities for the character-specific relative-rate parameters, $\sigma_i^2/c$, for each continuous character, under each of the specified prior densities. The boxes correspond to the 50% posterior credible interval, and the whiskers correspond to the 95% posterior credible interval.

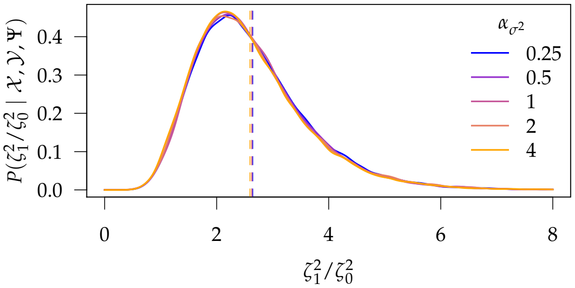


**Figure S11:** Marginal posterior probability densities of the relative-rate ratio, $\zeta_1^2/\zeta_0^1$, for each of the five priors on the character-specific relative-rate parameter, $\sigma^2/c$. The dashed horizontal lines are the posterior means for each of the five marginal posterior densities.

###### S.6.5.3 Discrete-character clock rate

We varied the prior on the rate of discrete-trait evolution, $\lambda$, such that the expected number of changes, $k$, varied over five values: $\mathbb{E}(k) \in \{1, 3, 5, 10, 20\}$. The prior densities on $\lambda$, and the corresponding marginal posterior densities, are depicted in Fig. S12. The induced priors on $k$, and the corresponding marginal posterior distributions, are depicted in Fig. S13. The marginal posterior densities for the relative-rate ratio, $\zeta_1^2/\zeta_0^1$, are depicted in Fig. S14.

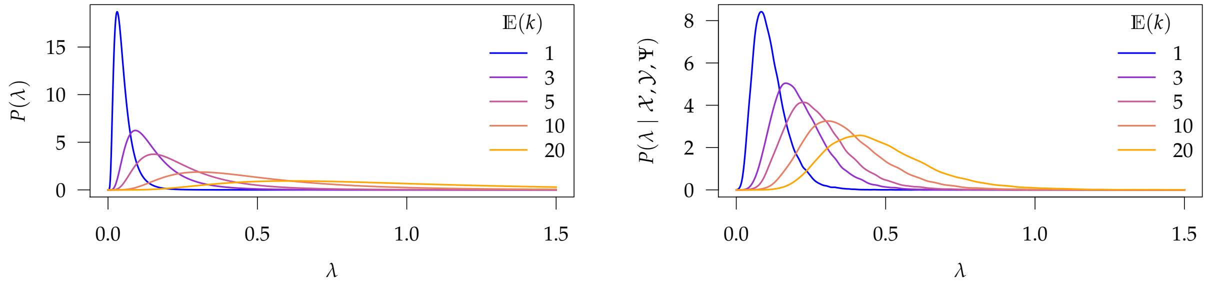


**Figure S12:** Prior sensitivity experiment for the rate of discrete-trait evolution, $\lambda$. Left panel: the five prior probability densities. Right panel: the five corresponding marginal posterior probability densities.

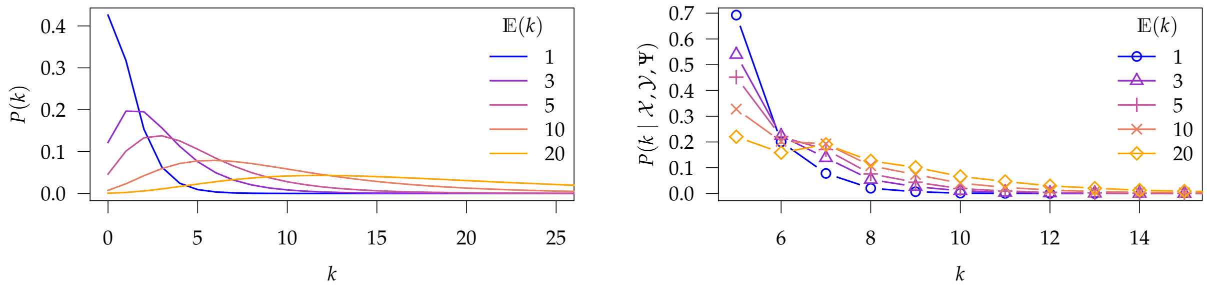


**Figure S13:** Prior sensitivity experiment for the induced prior on the expected number of discrete-character changes, $k$. Left panel: the five prior probability distributions. Right panel: the five corresponding marginal posterior probability distributions.

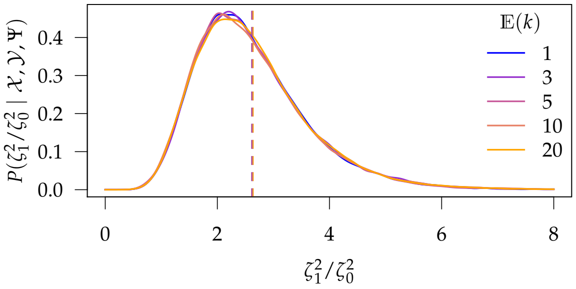


**Figure S14:** Marginal posterior densities of the relative-rate ratio, $\zeta_1^2/\zeta_0^1$, for each of the five priors on the rate of discrete-trait evolution, $\lambda$. The dashed horizontal lines are the posterior means for each of the five marginal posterior densities.

##### S.6.5.4 Background-rate variation

We varied the prior on the expected degree of background-rate variation, $\mathbb{E}(\nu)$, over three values: $\mathbb{E}(\nu) \in \{H \div 2, H, H \times 2\}$, where $\nu = H$ indicates that the 95% credible interval of the background rates covers one order of magnitude, $\nu = H \times 2$ indicates that they it covers two orders of magnitude, etc. These prior densities, and the corresponding marginal posterior densities, are depicted in Fig. S15. The marginal posterior densities of the relative-rate ratio, $\zeta_1^2/\zeta_0^1$, are depicted in Fig. S16. (Because $\boldsymbol{\zeta}^2/2$ is a vector of length 2, we plot the prior and posterior distributions of the first element, $\zeta_0^2/2$.)

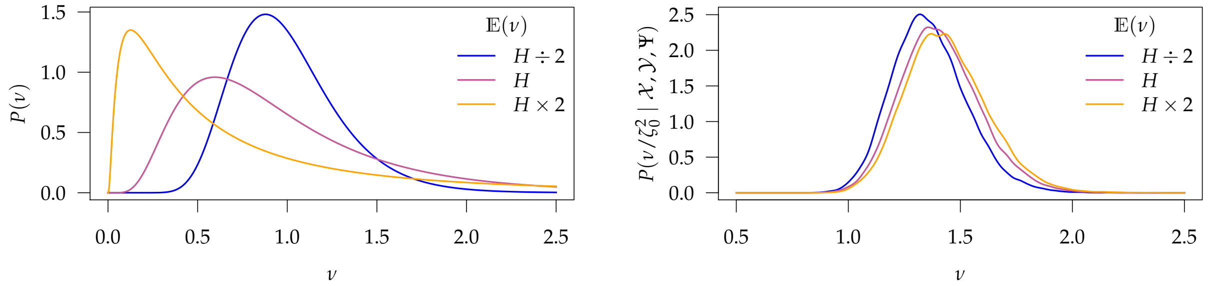


**Figure S15:** Prior sensitivity experiment for the expected degree of background-rate variation, $\mathbb{E}(\nu)$. Left panel: the three prior probability densities. Right panel: the three corresponding marginal posterior probability densities.

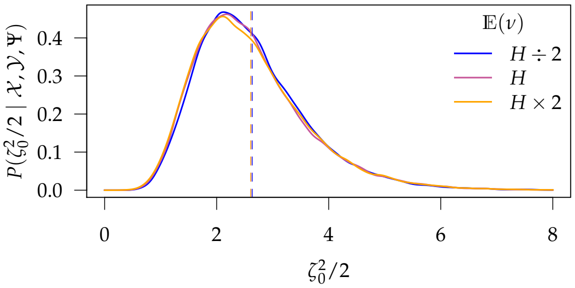


**Figure S16:** Marginal posterior densities of the relative-rate ratio, $\zeta_1^2/\zeta_0^1$, for each of the three priors on the expected degree of background-rate variation, $\mathbb{E}(\nu)$. The dashed horizontal lines are the posterior means for each of the three marginal posterior densities.

##### S.6.5.5 Correlation matrix

We varied the prior on the correlation matrix, $\eta$, over five values: $\eta \in \{1/2, 1, 2, 5, 10\}$. These priors, and the corresponding marginal posterior densities, are depicted in Figs. S17 and S18, respectively. The marginal posterior densities of the relative-rate ratio, $\zeta_1^2/\zeta_0^1$, are depicted in Fig. S19. (Because $R$ is a $c \times c$ matrix, we plot the distributions of the correlation between the $i^{\text{th}}$ and $j^{\text{th}}$ character, $\rho_{ij}$.)

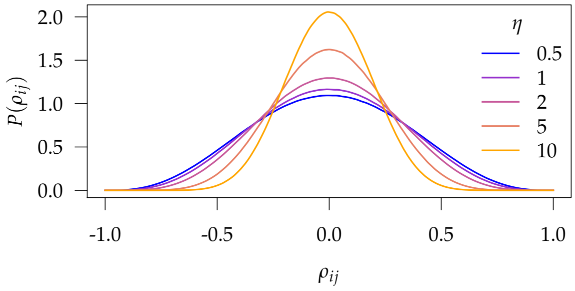


**Figure S17:** Marginal prior probability densities for pairwise correlation parameters between the $i^{\text{th}}$ and $j^{\text{th}}$ continuous characters, $\rho_{ij}$ for $c = 9$.

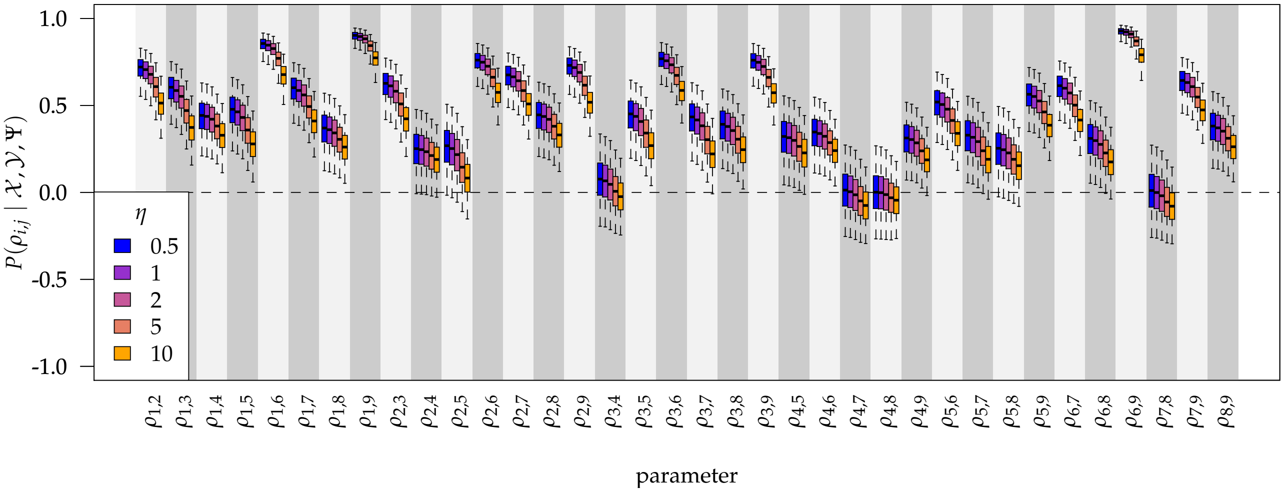


**Figure S18:** Boxplots of the marginal posterior probability densities for the pairwise correlation parameters, $\rho_{ij}$, for each pair of continuous characters, under each of the specified prior densities. The boxes correspond to the 50% posterior credible interval, and the whiskers correspond to the 95% posterior credible interval.

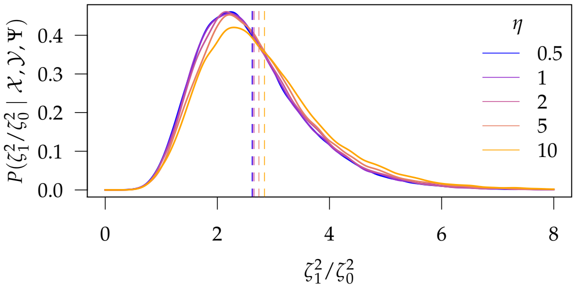


**Figure S19:** Marginal posterior probability densities of the relative-rate ratio, $\zeta_1^2/\zeta_0^1$, for each of the five priors on the correlation matrix, $R$. The dashed horizontal lines are the posterior means for each of the five marginal posterior densities.

#### S.7 Simulation Study

##### S.7.1 MCMC Analyses and Diagnosis

We performed two independent MCMC analyses for each unique dataset and model combination. We provide details of the specific proposals, proposal weights, and MCMC settings within the scripts included in our supplementary-data archive.

We diagnosed each analysis by computing the effective sample size and Geweke's diagnostic (Geweke et al. 1991) for each parameter using the R package `coda` (Plummer et al. 2006); we considered an analysis a failure if more than 5% of the diagnostics failed (ESS $< 500$, Geweke's $p$-value $< 0.05$). If an analysis failed these criteria, we re-ran it using double the number of MCMC generations, then re-assessed MCMC performance, as above. If an analysis failed a second time, we excluded it from our summaries of the statistical behavior. Finally, we combined samples from the two independent (and successful) MCMC analyses for down-stream summaries.

##### S.7.2 Statistical Behavior

We characterize the performance of the method under simulation using three statistics based on the posterior distribution: the *coverage probability*, which measures how often the true value of a parameter is contained within the posterior 95% credible region; the *accuracy*, which measures the proximity of the estimated parameter value (*i.e.*, the posterior mean) to the true parameter value; and the *precision*, which measures the width of the 95% credible region.

To compute the coverage probability, we compute the frequency of cases in which the true parameter value is contained within the 95% credible region (CR); for a well-behaved method, this frequency should be 95% (Huelsenbeck and Rannala 2004). For scalar parameters, the credible region is the credible interval, and it is straightforward to assess whether the true value is contained within the credible interval. For vector or matrix parameters, assessing parameter coverage is more complicated. For these parameters, we compute whether each element of the true vector or matrix is contained within its corresponding marginal posterior distribution, and then consider the true value covered with probability:

$$P(\text{covered}) = \frac{1}{n} \sum_{i=1}^{n} \mathbb{1}(\theta_i),$$

where $n$ is the number of elements of the parameter, and $\mathbb{1}(\theta_i)$ is an identity function that returns 1 when the $i^{\text{th}}$ element is covered, and otherwise returns 0. To compute the coverage probability for a set of analyses, we consider each analysis to be covered with probability $P(\text{covered})$, and then compute the fraction of analyses that are covered.

We measure the accuracy of the posterior-mean estimator of a parameter by computing the mean squared error for a given set of $n$ analyses:

$$\text{MSE} = \frac{1}{n} \sum_{i=1}^{n} d\left(\widehat{\theta}_i, \theta_i\right)^2,$$

where $d(\widehat{\theta}, \theta)$ is the "distance" between the true and estimated values of the $i^{\text{th}}$ parameter. For scalar parameters, we use the difference as the measure of distance, $d(\widehat{\theta}, \theta) = \widehat{\theta} - \theta$, and for multivariate parameters, we use the Euclidian distance between the true and posterior-mean estimate of the vector (or matrix) of parameters. We compute the mean of a multivariate posterior distribution as a combination of the mean of the marginal posterior distribution for each element. For example, for the correlation matrix $R$, we compute the posterior mean $\widehat{R}$, where the $ij^{\text{th}}$ element is the posterior mean of the correlation, $\widehat{\rho_{ij}}$. An additional complication of the MSE summary statistic is that values are difficult to compare across vectors and matrices of different dimensionality. To help "correct" for differences in dimensionality, we normalize the MSE for vectors and matrices by dividing by the average distance between many samples drawn from the corresponding prior distribution. In this case, the MSE actually measures the proximity of the estimated parameter value to the true parameter value compared to random draws from the corresponding prior distribution.

We measure the precision of the posterior distributions by computing the average width of the 95% posterior credible region across a set of analyses. Again, it is straightforward to measure the width of the credible region for scalars—it is just the average width of the credible intervals—but it is more difficult for vector and matrix parameters. For these parameters, we compute the Euclidian distance, $\delta$, between the posterior-mean estimate (as described above) and each of the sampled parameter values (*i.e.*, the set of parameter values in the MCMC sample), thereby providing a posterior distribution of $\delta$. Because this distance shrinks to zero as a sample grows close to the mean, we use the upper $95^{\text{th}}$ quantile as the measure of the width of the credible region. Again, this summary is difficult to compare across multivariate parameters of different dimensionality, so we normalize the value by the average $\delta$ between many samples drawn from each prior distribution.

In the following sections, we report the above summary statistics for the relevant model parameters: the state-dependent rate parameter, $\zeta_1^2$ (a scalar, since $\zeta_0^2 + \zeta_1^2 = 1$), the character-specific rate parameters, $\boldsymbol{\sigma}^2$ (a vector), the average background rate of continuous-character evolution, $\mu$ (a scalar), and the evolutionary correlation matrix, $R$ (a matrix). For each parameter, we report the behavior of the three statistics over the 100 simulations for each combination of $N \in \{25, 50, 100\}$, $c \in \{1, 2, 4, 8\}$, and $\zeta_1^2 / \zeta_0^2 \in \{1, 2, 4, 8\}$, first for the simulations under constant background-rates, and then for simulations under variable background-rates.

##### S.7.2.1 State-dependent rates

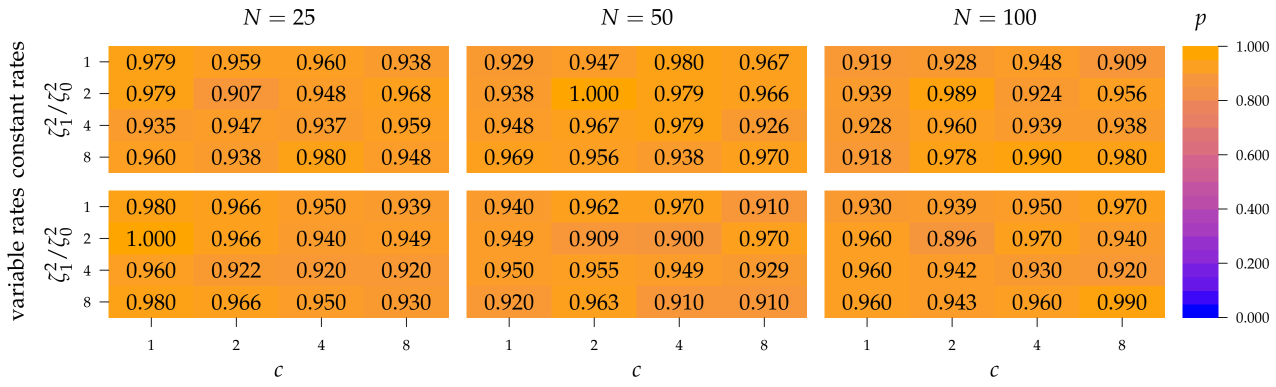


**Figure S20:** Coverage probability for the state-dependent rate parameters under constant background rates (top set of panels) and variable background rates (bottom set of panels). Each cell depicts the fraction of analyses in which the 95% CI contained the true parameter value for simulations under a specific combination of $N$, $c$, and $\zeta_1^2/\zeta_0^2$.

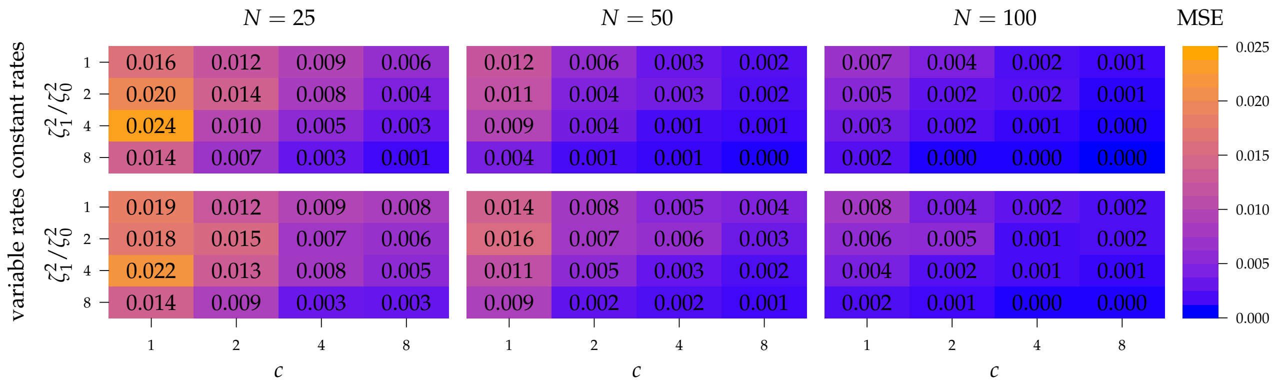


**Figure S21:** Mean squared error for the state-dependent rate parameters under constant background rates (top set of panels) and variable background rates (bottom set of panels). Each cell depicts the mean squared error for simulations under a specific combination of $N$, $c$, and $\zeta_1^2/\zeta_0^2$.

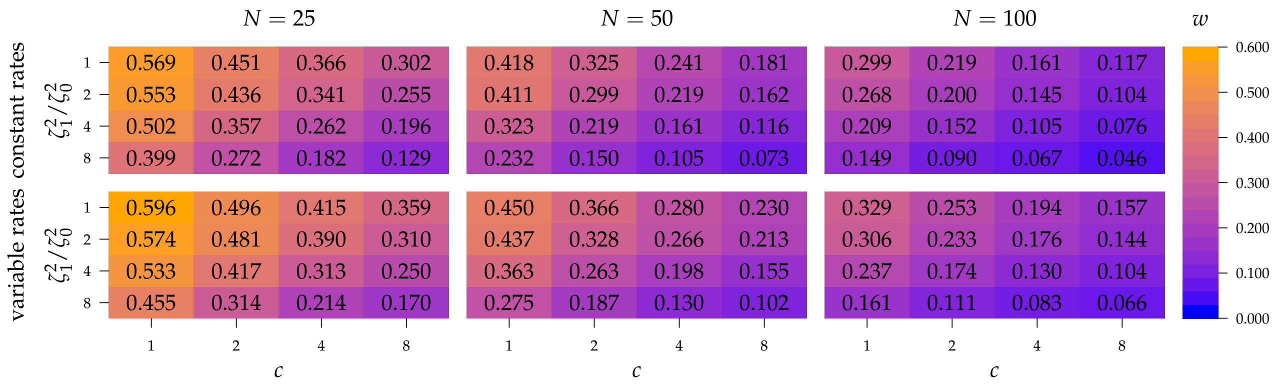


**Figure S22:** Precision for the state-dependent rate parameters under constant background rates (top set of panels) and variable background rates (bottom set of panels). Each cell depicts average "width" of the 95% credible interval for simulations under a specific combination of $N$, $c$, and $\zeta_1^2/\zeta_0^2$.

##### S.7.2.2 Character-specific rates

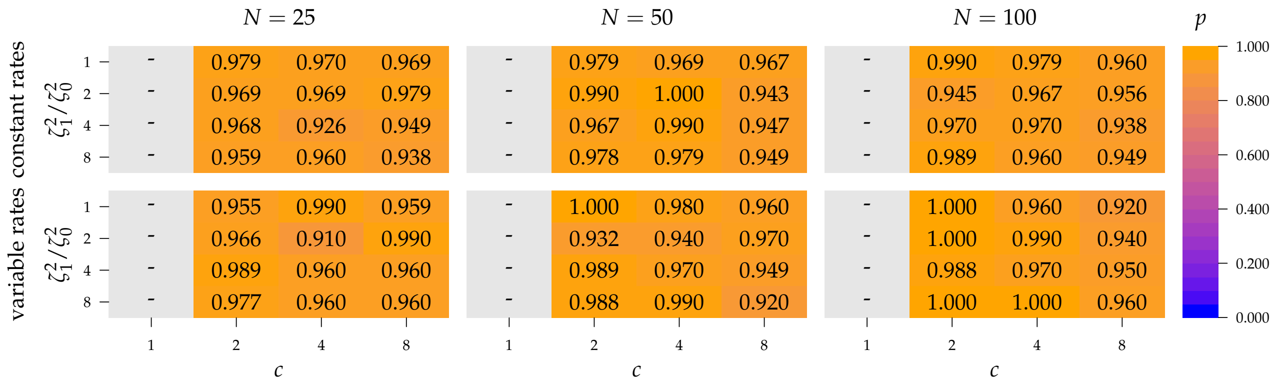


**Figure S23:** Coverage probability for the character-specific rate parameters (graphical conventions follow those described in Fig. S20).

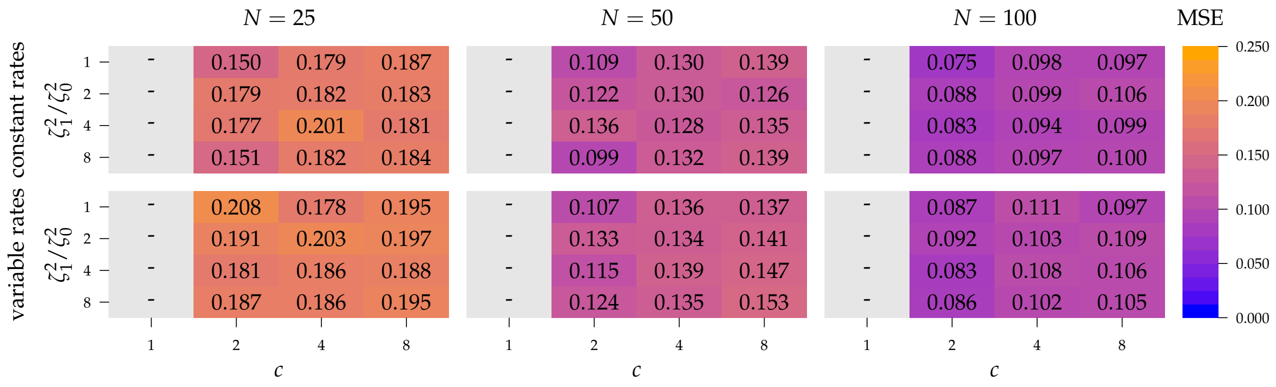


**Figure S24:** Mean squared error for the character-specific rate parameters (graphical conventions follow those described in Fig. S21).

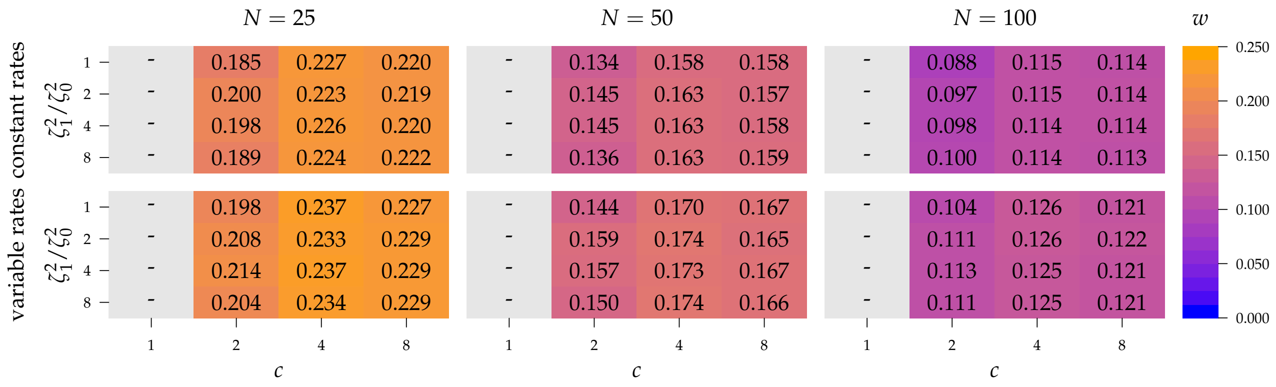


**Figure S25:** Precision for the character-specific rate parameters (graphical conventions follow those described in Fig. S22).

##### S.7.2.3 Background rates

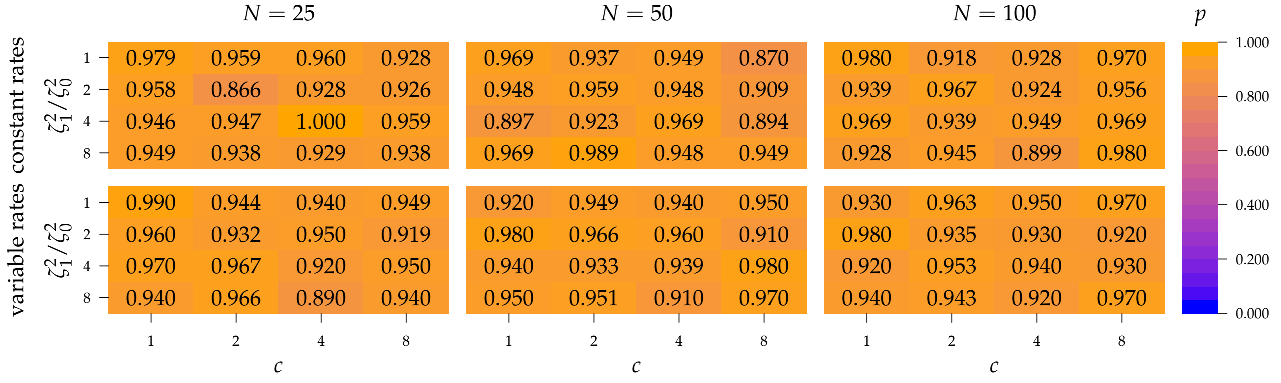


**Figure S26:** Coverage probability for the average background-rate parameter (graphical conventions follow those described in Fig. S20).

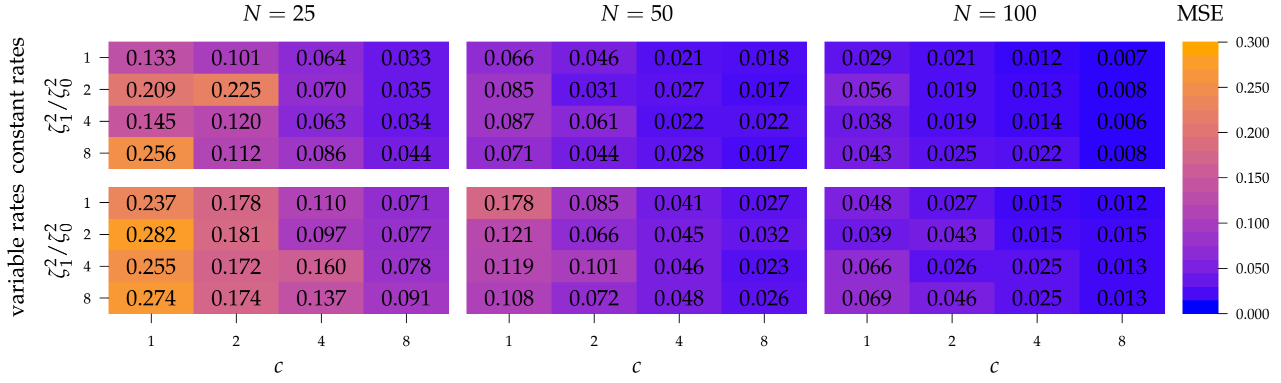


**Figure S27:** Mean squared error for the average background-rate parameter (graphical conventions follow those described in Fig. S21).

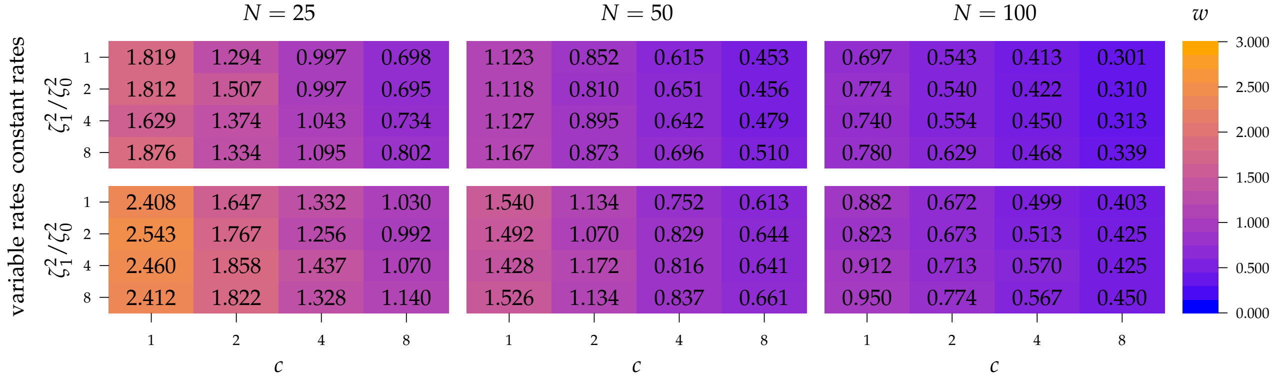


**Figure S28:** Precision for the average background-rate parameter (graphical conventions follow those described in Fig. S22).

##### S.7.2.4 Correlation matrix

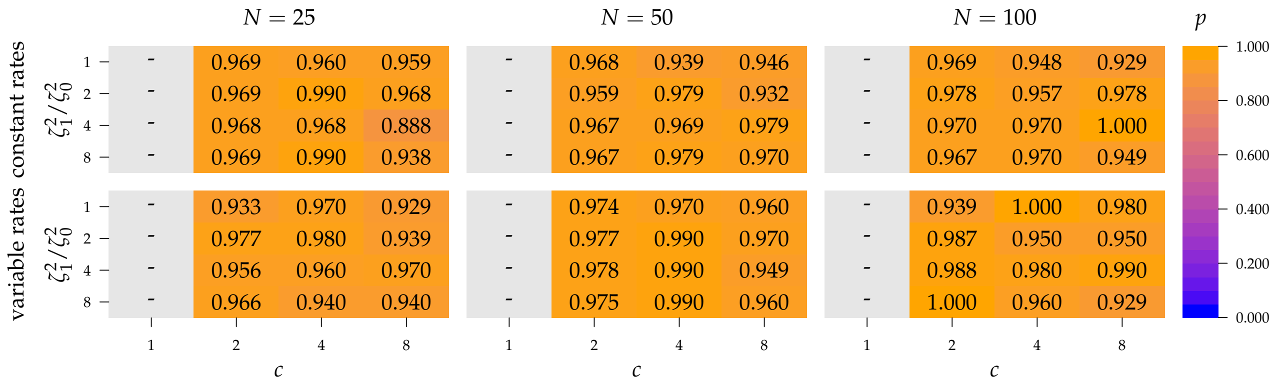


**Figure S29:** Coverage probability for the correlation matrix (graphical conventions follow those described in Fig. S20).

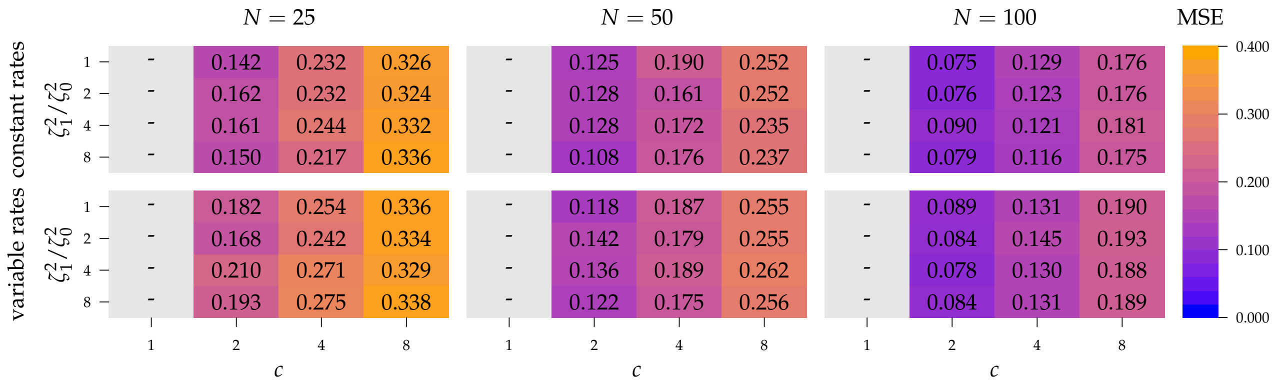


**Figure S30:** Mean squared error for the correlation matrix (graphical conventions follow those described in Fig. S21).

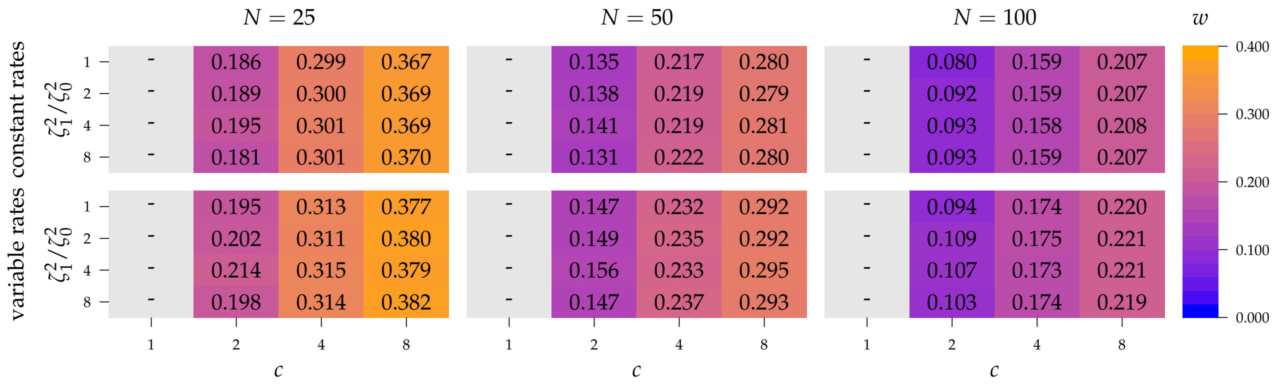


**Figure S31:** Precision for the correlation matrix (graphical conventions follow those described in Fig. S22).
